# Synaptomic analysis of dopaminergic inputs reveal hub synapses in the mouse striatum

**DOI:** 10.1101/2020.02.18.952978

**Authors:** Vincent Paget-Blanc, Marlene E. Pfeffer, Marie Pronot, Paul Lapios, Maria-Florencia Angelo, Roman Walle, Fabrice P. Cordelières, Florian Levet, Stéphane Claverol, Sabrina Lacomme, Melina Petrel, Christelle Martin, Vincent Pitard, Véronique Desmedt-Peyrusse, Thomas Biederer, David Perrais, Pierre Trifilieff, Etienne Herzog

**Author notes:** These authors contributed equally to the work (Vincent Paget-Blanc, Marlene E. Pfeffer, Marie Pronot). Correspondence should be addressed to Etienne HERZOG, Université de Bordeaux, Interdisciplinary Institute for NeuroScience - UMR 5297, Centre Broca Nouvelle-Aquitaine, CS 61292 Case 130, 146 rue Léo Saignat, 33076 Bordeaux Cedex (FRANCE).

## Abstract

Dopamine is a monoamine involved in reward processing and motor control. Volume transmission is thought to be the mechanism by which monoamines modulate effector systems at glutamate and GABA synapses. Hence, dopamine synapses are scarcely described. We applied fluorescence activated synaptosome sorting to explore the features of the dopaminergic synaptome. We provide the proteome of striatal dopaminergic synapses with 57 proteins specifically enriched. Beyond canonical markers of dopamine neurotransmission (Th, Slc6a3/DAT, Slc18a2/VMAT2), we validated 6 proteins belonging to pre- and postsynaptic sides (Cpne7, Apba1/Mint1, Cadps2, Cadm2/SynCAM 2, Stx4 and Mgll). Moreover, dopaminergic varicosities adhere to both a post-synapse with cognate receptors and glutamatergic, GABAergic or cholinergic synapses in structures we named dopaminergic “hub synapses”. Markers of presynaptic vesicles and active zone, post-synaptic density and spine apparatus, are significantly increased upon association with dopamine inputs in hubs. Thus neuromodulation frequently operates from hub synapses affecting associated synapses and is not solely dependent on volume transmission. Finally, FASS provides a new framework for the exploration of dopaminergic synapses and more generally for discrete synapse populations ex-vivo.

**Highlights:**

1. A first proteome of dopaminergic synapses in the striatum
2. Striatal dopaminergic synaptosomes display post-synaptic cognate receptors
3. Dopaminergic projections build hub synapses with excitatory, inhibitory, and cholinergic projections.
4. Cortico-striatal synaptic scaffolds are strengthened upon association in hub synapses.

## INTRODUCTION

Since the 1950’s with the first ultrastructural characterization of the synapse in the central nervous system (Gray, 1959) a wide variety of synapse types have been described based on morphological criteria (Harris). The archetypal synapse type extensively studied is the asymmetric excitatory synapse on dendritic spines (Gray, 1959) whose ultrastructure is easily identifiable in the tissue and routinely studied *in vitro* using primary neuronal cultures (Banker & Cowan, 1977). Alternatively, symmetric synapses are mostly inhibitory or modulatory. They do not display post-synaptic densities and are more difficult to identify *in situ* (Descarries *et al*, 1996; Moss & Bolam, 2008). Moreover, many types of synaptic organizations are not abundant enough and/or accessible in *in vitro* models. These limitations hinder our understanding of neuronal network functioning.

While glutamate and GABA (Gamma-Amino Butyric Acid) neurotransmissions drive point to point information locally, modulatory neurotransmitters pace regional activity through volume transmission in the neuropil (Agnati *et al*, 1995; Greengard, 2001). Dopamine transmission is a major neuro-modulatory system involved in several functions such as movement initiation, reward prediction error and incentive processes, notably by its projections onto spiny projection neurons (SPNs) of the striatum (Kreitzer, 2009). Dopamine signalling is presumed to modulate glutamate transmission onto SPNs through release of dopamine mainly from varicosities devoid of synaptic differentiation while a minority forms synapses onto SPN spines or dendrites, as well as presynapses (Descarries *et al*, 1996; Moss & Bolam, 2008; Bamford *et al*, 2004). However, recent work challenges the model of volume dopamine transmission by providing intriguing evidence for local point-to-point signalling. In fact, optophysiology approaches revealed rapid and local transmission at dopaminergic projections to the striatum (Yagishita *et al*, 2014; Howe & Dombeck, 2016; Pereira *et al*, 2016), which is in accordance with the requirement for synaptic vesicle release machinery for fast dopamine release at striatal varicosities (Liu *et al*, 2018). Moreover, the distribution of varicosities in the striatal neuropil appears biased toward proximity with the surrounding effector synapses (Moss & Bolam, 2008), and dopamine receptors interact physically and functionally with glutamate and GABA receptors (Liu *et al*, 2000; Cahill *et al*, 2014; Ladépêche *et al*, 2013; Cepeda & Levine, 2012), suggesting a tight coupling between dopamine and effector transmissions.

In the present work we aimed to unravel the cellular and molecular synaptome of single projection pathways (Zhu *et al*, 2018). This can critically complement current connectomic approaches using optophysiology and tracing methods, which are limited in terms of molecular analysis of specific synapses at play in a given circuit (Schreiner *et al*, 2016). To that end, we established a workflow combining fluorescence tracing of the dopaminergic pathway, fluorescence activated synaptosome sorting and an array of semi-quantitative analysis methods ranging from conventional immunofluorescence characterization to mass spectrometry-based proteomics. With this approach we provide the first ex-vivo model to thoroughly analyse the cellular and molecular organisation of dopaminergic synapses from mouse striatum. This new model unravels the existence of a physical coupling between dopaminergic and effector synapses in a complex we name “hub synapses”. Synaptic hubs may represent key units in the modulatory action of dopamine on glutamate and GABA signalling.

## RESULTS

### Fluorescence activated synaptosome sorting (FASS) enrichment of dopaminergic synaptosomes reveals synaptic hub structures

Here we labelled the Dopaminergic projection onto the striatum through stereotaxic injection of a viral vector carrying Cre-dependent EGFP (Oh *et al*, 2014) in the midbrain of Dopamine Transporter promoter (DAT)-Cre transgenic mice (Turiault *et al*, 2007) (Figure 1A1-2; for increased yields we also used mNeonGreen as a fluorescence reporter see Figure S1). We miniaturized the classical sucrose synaptosome fractionation to 1.5ml tubes (Whittaker, 1993; De-Smedt-Peyrusse *et al*, 2018) (Figure 1A3). Fluorescence activated synaptosome sorting (FASS) (Biesemann *et al*, 2014; Luquet *et al*, 2017) applied to this sample allowed recovering up to 35 million synaptosomes for immunoblot and analysis of protein content by mass spectrometry (Figure 1A4-6). In addition, we established the immobilisation of particles on glass coverslips to analyse them through quantitative immunofluorescence, super-resolution STED microscopy and electron microscopy (Figure 1A5-6). To validate our labelling approach we performed a complete subcellular fractionation of the samples dissected from the striatum and measured the amount of tyrosine hydroxylase (Th), a soluble protein of dopaminergic presynapses that catalyses the limiting step for dopamine synthesis (Lamouroux *et al*, 1982), and the soluble reporter EGFP. Subcellular fractions were probed using a semi-automatic capillary immunoblot system producing electropherograms (Figure 1B) or membrane-like band patterns (Figure 1C). Quality controls of the fractionation shows the enrichment of synaptophysin (Syp) in synaptosomes (SYN) and crude synaptic vesicle (LP2) fractions while the plasma membrane glutamate transporter GLAST (Slc1a3/GLAST) is enriched in synaptic plasma membranes (SPM). We confirm the concentration of Th and EGFP signals in soluble fractions of synaptosomes (LS1 and LS2) while soluble proteins of the homogenate display weak Th and EGFP signals (S2; Figure 1CD). Hence, most of the soluble protein content of dopaminergic axons is engulfed in synaptosomal membranes and available for discrimination in our FASS procedure. Our gating strategy was adapted from previous work (Biesemann *et al*, 2014) in order to avoid sorting aggregated particles (Figure S1) and detect EGFP particles specifically and with high probability (Figure 1E). Synaptosomes from DAT-EGFP tracings contained on average 3.9% singlet EGFP positive synaptosomes (3.9% ± 0.52; range 2-6 % n = 9; Figure 1F) upon reanalysis in the cell sorter after FASS EGFP synaptosomes represented around 50% of the total (49% ± 2.3%; N = 9 sorts; Figure 1F). EGFP-particles in the synaptosome samples were depleted accordingly (Figure 1F). We first validated these sorts using capillary electrophoresis-based immunoblots. As expected, Th and the dopamine transporter (Slc6a3/DAT) display a steep enrichment after DAT-EGFP FASS. In contrast, GLAST is strongly depleted while the glutamate receptor (Gria1/GluA1) or the synaptic active zone protein Munc18 (Stxbp1) are only slightly reduced (Figure 1G). We then explored FASS samples using transmitted electron microscopy (Figure 1H-L). We easily identified synaptosome profiles with resealed presynaptic elements (Figure 1H) and in some cases a clear adhesion with a post-synaptic membrane (Figure 1I). Surprisingly, we also identified a significant proportion of profiles displaying several presynapses organized around possible postsynaptic membranes (Figure 1J-L). In many occurrences, the synaptosomes were cut with an angle that prevented clear identification of all synaptic elements (Figure 1L). On another example, we found 2 distinct presynapses, one electron dense terminal with many synaptic vesicles adhering to a presynaptic element with fewer vesicles and to another compartment that could be dendritic (Figure 1J). Finally, a post-synaptic element displayed adhesion to 3 different “boutons”, one of them displaying a clearer background and less vesicles (Figure 1K). These complex micro-structures were preserved even though our procedure exposed the tissue to shearing forces during homogenization. Moreover, additional shearing forces prone to collapse aggregates are applied to particles in the nozzle of the cell sorter (see workflow in Figure 1A). This suggests the existence of a specific adhesive interaction between partners of these structures that we named “hub synapses”. We then pursued the characterization of dopaminergic hub synapses to identify their molecular nature.

**Figure 1:**
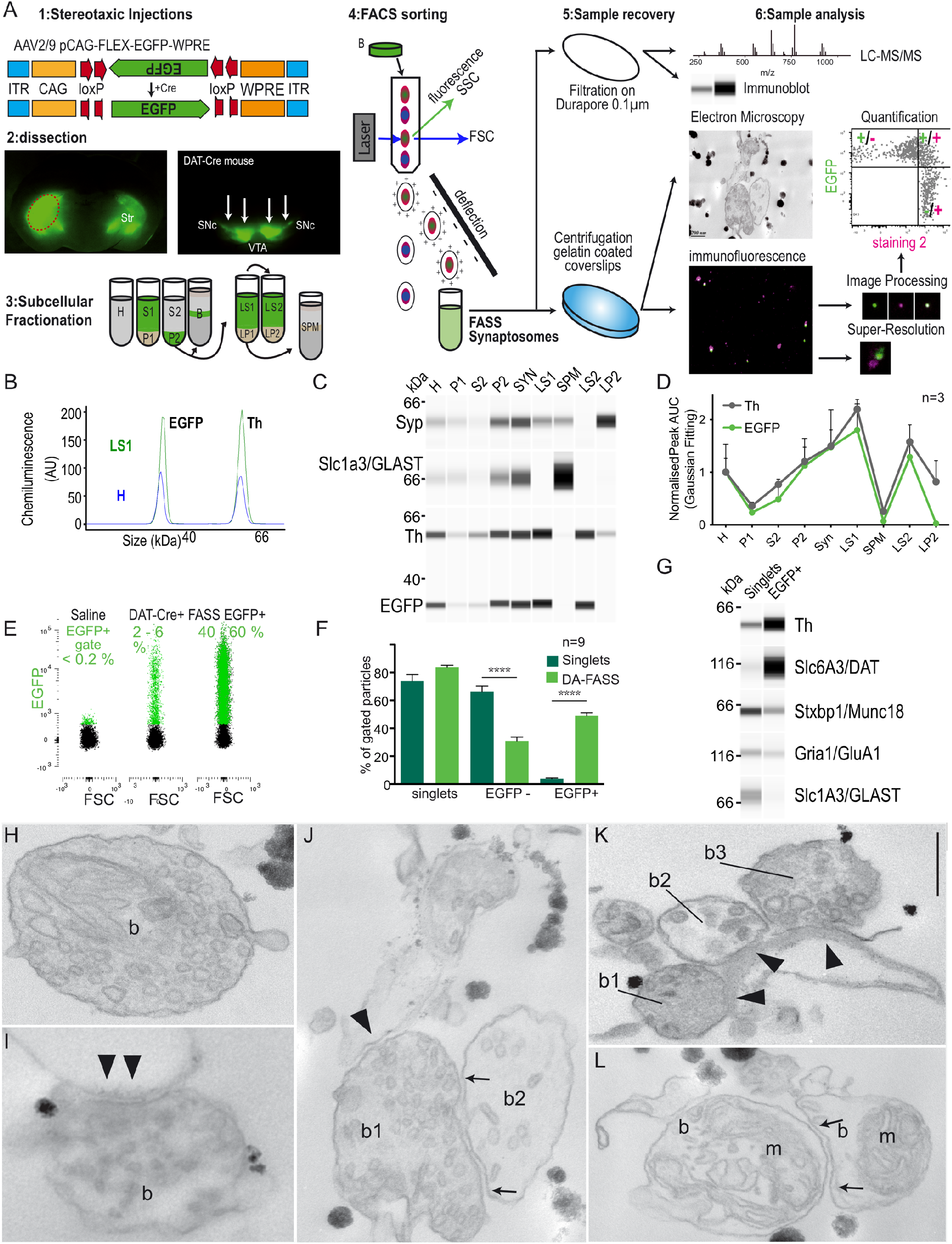
single projection Fluorescence Activated Synaptosome Sorting (FASS) isolates dopaminergic hub synaptosomes. **(A)** Workflow of DAT-cre/AAV-EGFP based synaptosome sorting and analysis. (1) DAT-Cre+ mice stereotaxically injected with a Cre-dependent AAV expressing EGFP or mNeongreen (Figure S1) in the Substantia Nigra pars compacta and the Ventral Tegmental Area. (2) Dissection of brightest fluorescent part of Striatum (Str) (red-dashed circle). (3) Synaptosome subcellular fractionation on a discontinuous sucrose gradient and (4) Fluorescence Activated Synaptosome Sorting (FASS). (5) Collection by filtration or centrifugation on glass coverslips. (6) FASS sample analysis by mass spectrometry, immunoblot, electron microscopy, conventional and super-resolved immunofluorescence. **(B-D)** Analysis of subcellular fractionation through capillary electrophoresis based immunoblot. (B) EGFP and TH fitted chemiluminescence peaks for H (blue) and LS1 (green) fractions. (C) Representative chemiluminescence bands of Synaptophysin1, Slc1a3/GLAST, Th and EGFP. (D) relative integrated intensity for Th (grey) and GFP (green) for each subcellular fraction (H to LP2; n=3 complete fractionations). Note that most Th and EGFP proteins are concentrated in synaptosomes (P2 and Syn) and get released with soluble proteins upon osmotic lysis (LS1 and LS2), all data are mean ±SEM. **(E)** Analysis of synaptosomes through FASS sorting. (left) Saline synaptosomes set the level of autofluorescence. Gating was set to have 0-0.2% of particles within the EGFP+ range in negative controls. (middle) Singlets DAT-Cre+ synaptosome before sorting show 2-6% of EGFP+ particles. (right) FASS EGFP+ synaptosomes reanalysed in the sorter consist of 40-60% of EGFP+ particles. See Figure S1 for detailed gating strategy **(F)** Average DAT-cre/EGFP+ Singlets and FASS results after sorting. Singlets (dark green) and FASS (green) samples for the different gates: singlets, EGFP-, EGFP+. Note the steep increase in EGFP+ particles and significant decrease in EGFP-contaminants through the FASS process. n=9 sorts, all data are mean ±SEM. ****p < 0.001, Two-way MD ANOVA, post-hoc Šídák. Complete statistics are available in supplementary table 2**. (G)** Immunoblot analysis of DA-FASS samples. Compared to unsorted singlets, EGFP+ FASS samples display strong increase in Th and Slc6a3/DAT signals, a moderate reduction in the presynaptic marker Stxbp1/Munc18 or the post-synaptic glutamate receptor Gria1/GluA1, but a steep decrease in the astrocytic plasma membrane transporter for glutamate Slc1a3/GLAST. **(H-L)** Electron micrographs of sorted synaptosomes. **(H-I)** Typical synaptosomes displaying synaptic vesicles (SV) rich bouton (b) and synaptic contact with an opened postsynaptic membrane (arrowheads in I only). **(J)** Example of complex synaptosome that we named “hub synapse” displaying a SV-rich bouton (b1) contacting a postsynaptic membrane (arrowheads) and a second bouton (arrows) less populated with SVs (b2). **(K)** Synaptic hub displaying 3 distinct presynaptic profiles (b1, b2, and b3) contacting a postsynaptic membrane (arrowheads). Note the middle bouton (b2) less populated with SVs. Scale bar, 200 nm. Unspecific electron dense precipitates result from the embedding of synaptosomes on gelatin chrome-alum coating. **(L)** example of a synaptic hub structure cut through a plan that is not optimal to identify all synaptic elements.

### DAT-EGFP FASS synaptosomes display pre- and post-synaptic features of dopaminergic synapses

What is the molecular makeup of these biochemically isolated hub synapses? We addressed this by comparing both unsorted singlets and FASS-sorted synaptosomes using immunofluorescence detection of dopaminergic markers. Individual particles were quantified and results were plotted according to their intensity in both channels. Quadrant gates were defined to split positives and negatives for each label (see Figure 1A6). The top 2 quadrants are EGFP+ particles and percentages of particles are displayed in each quadrant. TH-positive, EGFP-labelled synaptosomes population rose from 54% of the total before sort to 82% after sort. The number of labelled particles per field of view was increased around 5-fold over sorting (Figure 2ABC). STED imaging revealed that TH signals were highly colocalized with EGFP (Figure 2D). Similarly, we found a strong co-localization with anti-DAT (Dopamine Transporter) signal. Yet, a significant number of DAT+/EGFP-observed may correspond to extra synaptic axonal pieces that did not retain the cytosolic EGFP (Figure S2AB). As expected from our immunoblot analysis (Fig 1G), the marker Slc1a3/GLAST that labels astrocyte membranes was not significantly associated to the EGFP-labelled synaptosomes (Figure S2CD). This data further confirms that DAT-EGFP labelled synaptosomes display genuine dopaminergic synaptic markers and are strongly enriched through FASS.

**Figure 2:**
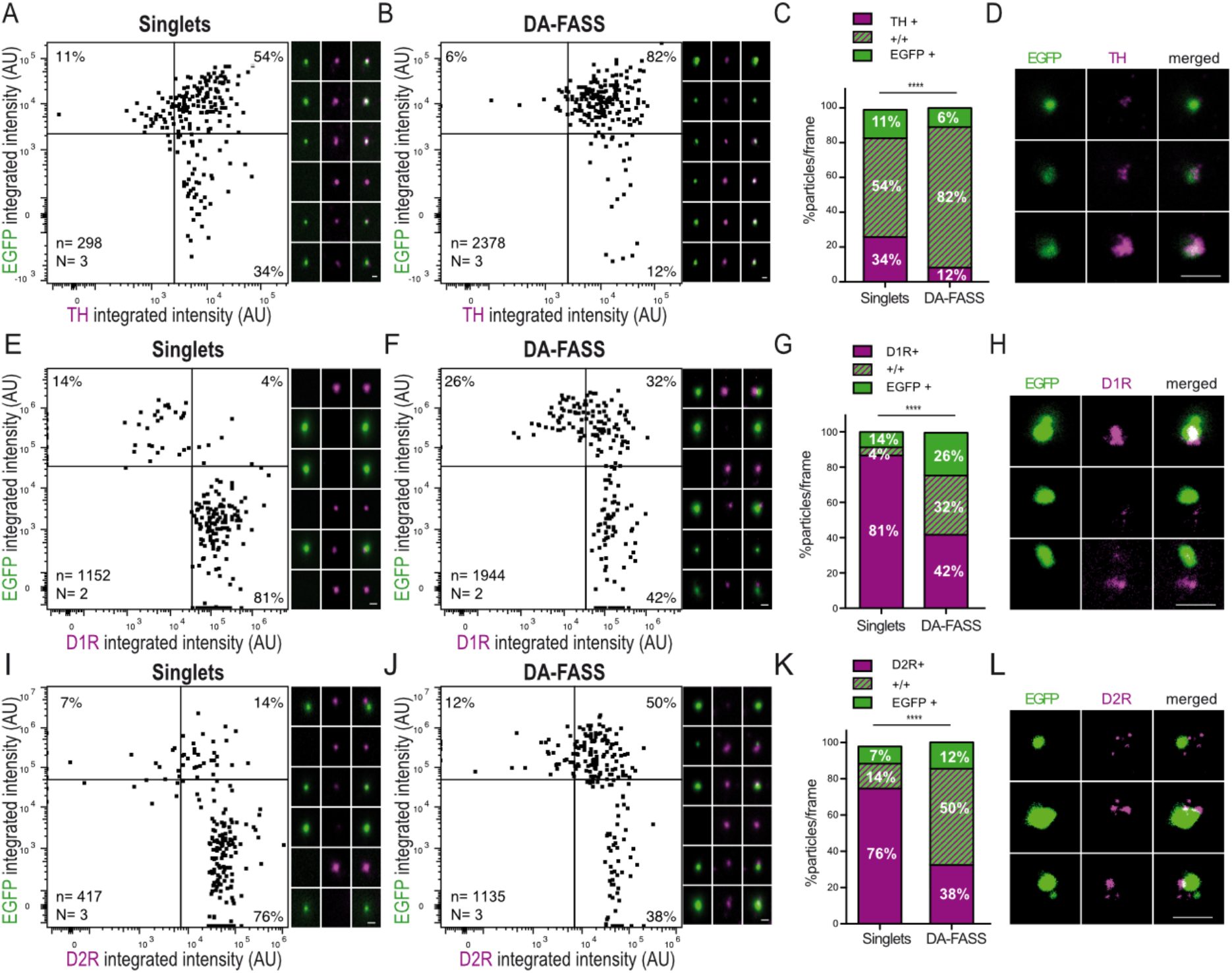
Immunofluorescence analysis of DA FASS synaptosomes reveal the enrichment for pre- and post-synaptic dopaminergic markers. **(A-B)** Dot plots (left) and epifluorescence images (right) of a representative sample of synaptosome populations (singlets in A and EGFP+ FASS in B) labeled with anti-TH (x-axis) and anti-EGFP (y-axis). Double positive particles population increases from 54% (A) to 82% (B). **(C)** Analysis of staining as in (A) and (B) showing particle proportions per frame. mean, interaction ****p < 0.001. Two-way MD ANOVA. **(D)** STED images of EGFP (green) and TH (magenta) labelled synaptosomes. Note the nearly perfect colocalizations. **(E-F)** Same as A-B for EGFP and D1 dopamine receptors. Double labeled populations increase from 4% (E) to 32% (F). Note that D1R positives represent about 55% of EGFP+ (dopaminergic) particles, while D1R+/EGFP-may represent extra-synaptic receptors on contaminants of the FASS sample. **(G)** Proportion of differently stained particles per frame, mean, interaction ****p < 0.001. Two-way MD ANOVA. **(H)** STED microscopy detects D1 receptor clusters (magenta) apposed to the EGFP+ synaptosomes (green). **(I-J)** Same as A-B for anti-EGFP and anti-D2 dopamine receptors. Double positive synaptosome representation increases from 14% (I) to 50% (J). **(K)** Proportion of differently stained particles per frame. mean, interaction ****p < 0.001. Two-way MD ANOVA. **(L)** STED images display D2R (magenta) patches apposed to EGFP (green). For all panels, scale bar = 1 μm. See extra immunofluorescence analysis in Figure S2 and complete statistics in supplementary table 2.

We then explored the co-segregation of dopamine receptors type 1 and −2 (D1R, D2R) together with DAT-EGFP labelled varicosities. D1R co-enriched 8-fold with DA-FASS (from 4% to 32%; Figure 2EF top right quadrants), while extra-synaptic (EGFP-) D1R labelled particles depleted 2-fold (from 81% to 42%; Figure 2EF bottom right quadrants). 55% of DAT-EGFP synaptosomes were labelled for D1R (EGFP+/D1R-26%, EGFP+/D1R+ 32%; Figure 2E-G upper quadrants). Anti-D1R displayed patches of staining apposed to EGFP particles (Figure 2H). D2R labels were found on more than 80% of DAT-EGFP dopaminergic synaptosomes and co-enriched massively with EGFP (EGFP+/D2R-12%, EGFP+/D2R+ 50%; Figure 2I-K upper quadrants). Extra synaptic (EGFP-) D2 receptors were depleted more than 2-fold over DA-FASS (EGFP-/D2R+ 76% in singlets vs 38% in FASS samples; Figure 2I-K lower right quadrants). D2R are auto- and hetero-receptors (Sesack *et al*, 1994), therefore, D2R found closely or more distantly apposed to EGFP labelled synaptosomes (Figure 2L) are likely to correspond to pre- and postsynaptic D2R, respectively. Altogether our data supports that nearly all dopaminergic synaptosomes carry a post-synaptic element equipped with cognate receptors.

### Label-free semi-quantitative proteomics reveals 57 proteins highly enriched at DA-FASS synaptosomes

Upon validation of our DA-FASS approach, we aimed at generating a set of samples for detection and quantification of the protein composition of dopaminergic synaptosomes through mass spectrometric (MS). We accumulated 35×10^6^ particles in three pairs of samples. Unsorted singlets representing the bulk synaptosome preparation were used as control samples and DA-FASS representing our dopaminergic sample. Protein identification and quantification yielded 3824 proteins identified with 1 peptide or more among which 2653 proteins could be quantified with at least 2 peptides. Proteins quantified with 2 peptides or more, displaying a reduction greater than 25% or an increase greater than 150% compared to the control, and with an ANOVA p value below 0.05, were considered significantly different from the unsorted control sample. Under these criteria, 63 proteins are significantly depleted upon sorting while 57 others appear significantly enriched (Figure 3A, Table S3).

**Figure 3.**
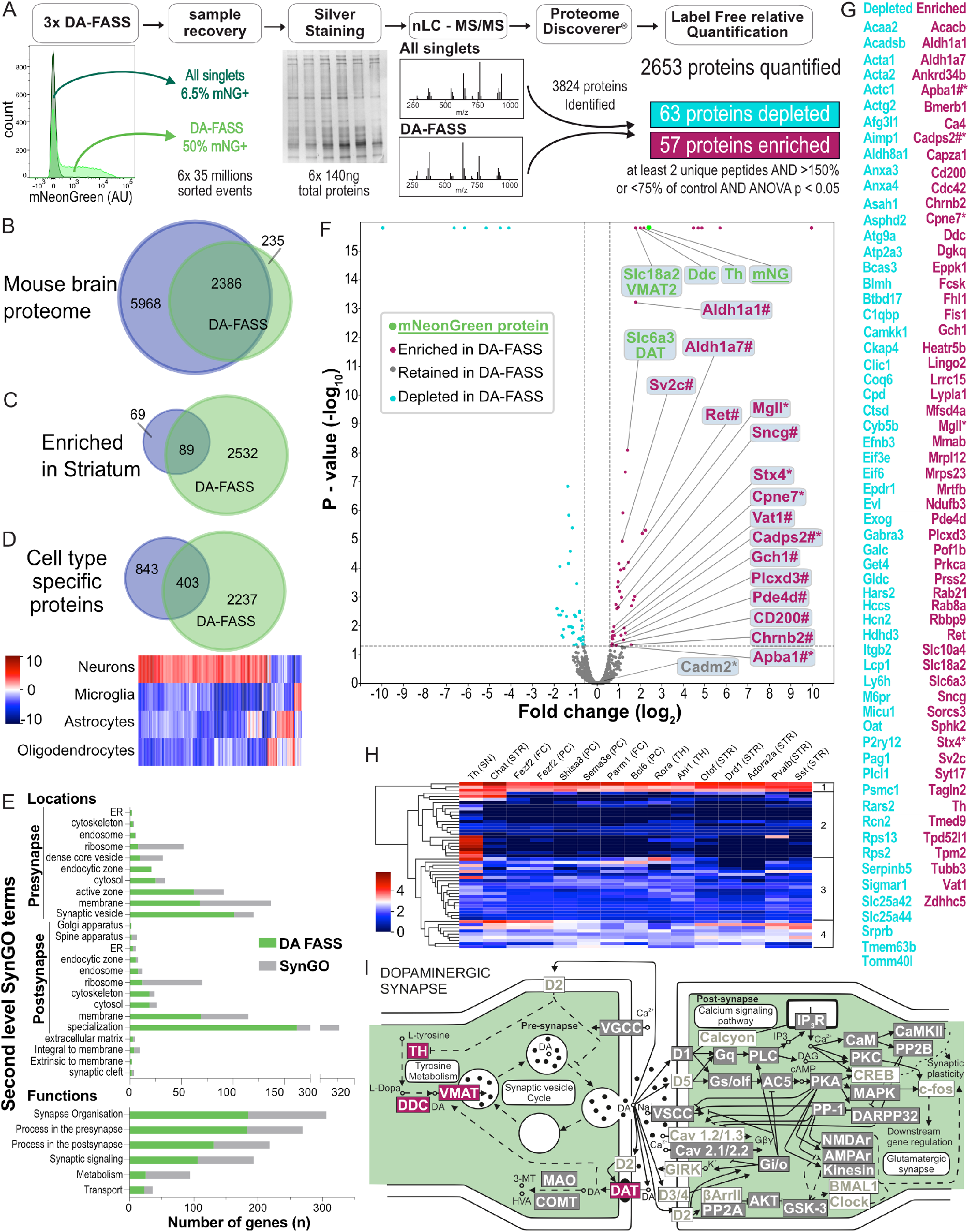
Comparative proteomic analysis of singlets and FASS purified DAT-mNeongreen positive synaptosomes. **(A)** Workflow of DAT-cre/AAV-mNeongreen singlets and FASS purified synaptosomes quantitative proteomic analysis. Following sample recovery samples were analyzed by silver staining. Protein content was normalized to 140ng for each replicate and loaded onto SDS-PAGE gel followed by tryptic digestion. Proteins of both samples were analyzed by high-resolution tandem MS. Of 2653 quantified proteins, 63 were depleted while 57 were enriched (Supplementary Table S3). **(B-D)** Venn diagrams representing the comparison of DA-FASS proteome with the mouse brain proteome resource (Sharma *et al*, 2015) as a whole (B), only enriched in the striatum (C) and cell type specific (D). 2386 DA-FASS proteins were present in the mouse brain proteome database (B). 89 out of the 158 striatal protein enriched from the mouse brain proteome database were found in our sample (C). (D)(top) From the 1246 proteins identified to be specific of a given cell type in cultures, 403 overlap with our synaptosome samples. (bottom) Heatmap showing cultured cell type protein abundance (as expressed in Sharma *et al*, 2015) of the list of overlapping proteins. Note that our striatal synaptosome proteome is highly biased toward neuronal specific proteins. **(E)** Subcellular localizations and function of the 2653 proteins quantified in the DA-FASS sample classified by second level SynGO terms (Koopmans *et al*, 2019). **(F)** Volcano plot of DA-FASS proteins. Values are plotted for each protein fold change versus their *P* value (on logarithmic scales). Thresholds are set at ± 1.5-fold change and p<0,05. Proteins are colored by subclass of canonical (green) enriched (red), depleted (cyan), and retained (grey) in the DA-FASS sample. # for proteins previously described as playing a role in dopamine signaling are annotated. * for targets selected for further experimental validations. **(G)** Complete list of depleted and enriched DA-FASS proteins. **(H)** Heatmap showing cell type specific mRNA abundance of the enriched DA-FASS proteins in striatal neurons (STR) or afferent cells to the striatum (Substantia Nigra, SN; Thalamus, TH; Frontal Cortex, FC; Posterior Cortex, PC) (data from DropViz; (Saunders *et al*, 2018)). Hierarchical clustering display 4 major clusters relating to the selectivity of mRNA expression. A more detailed heatmap can be found in supplementary figure 3 (S3). **(I)** Schematic of the molecular organization of a dopaminergic synapse (Adapted from the database KEGG). Enriched proteins from our DA FASS sample are in red, retained in grey, and absent in white. Gene names for each protein class can be found in supplementary table 5 with absent ones greyed out (Table S5).

We first compared our 2653 proteins dataset to the previous broad survey of mouse brain proteins produced by Sharma and colleagues (Sharma *et al*, 2015). 90% of our dataset is common to the global mouse brain proteome and of the 158 proteins significantly enriched in the bulk dissection of the striatum, 89 are represented in our synaptosome samples which is consistent with the selectivity of our subcellular fractionation (Figure 3BC, Table S4). Among the 1246 proteins identified to be specific of a given cell type in cultures we see an overlap with 403 of our synaptosome samples. A heatmap analysis of these shows the main neuronal origin of our synaptosome sample (Figure 3D, Table S4, Figure S2). We then performed a comparison of our proteome with the database of synaptic gene ontologies (SynGO)(Koopmans *et al*, 2019). Among 2653 genes of our proteome, we identified 684 genes documented in SynGO. These cover all localizations and functions reported in the second level of SynGO terms. For most SynGO terms, our gene set covers a majority of previously identified synaptic genes of the category (Figure 3E).

DA-FASS significantly depleted 63 proteins in our samples without a clear gene ontology signature. Beyond 57 proteins highly enriched during DA-FASS procedure, we could identify the strong enrichment of our reporter protein mNeonGreen (12 unique peptides, 5.12-fold increase, p value=1.6×10^−16^; Figure 3F, Table S3). mNeonGreen enrichment thus represents the target enrichment value for the most specific dopaminergic proteins. In line with this, the major canonical proteins involved in dopamine metabolism (Th; Ddc: DOPA decarboxylase; Slc18a2/VMAT2: Vesicular Monoamine Transporter type 2) show similar enrichment values. Slc6a3/DAT displays a slightly lower enrichment that may be explained by the loss of DAT proteins present on the axon shaft between varicosities (Figure 3F and S2)(Rahbek-Clemmensen *et al*, 2017). Of note, 14 proteins quantified with only 2 peptides displayed enrichment scores much higher than the 5-fold increase of our reporter. The extent of enrichment is possibly distorted by a weak detection in MS/MS. We identified a set of 12 proteins that were previously shown to be important for dopamine signaling and are seen enriched in our dataset (Figure 3FG marked with a #). Based on these elements, we chose a set of 6 proteins involved in membrane traffic, cell adhesion and neurotransmission to focus our validation efforts (Fig 3FG marked with a *). We probed the cell type expression pattern of the 57 enriched proteins with DropViz single cell RNA sequence database. For this analysis we focused on afferent and efferent neurons to the mouse striatum (Saunders *et al*, 2018). This meta-analysis provides hints regarding the identity of neurons expressing the enriched markers. It defines 4 clusters of gene expression, from ubiquitous expression to expression restricted to Th neurons of the midbrain. This analysis further suggests that some of the DA-FASS enriched proteins belong to various partners of the synaptic hubs (Figure 3H and S3). Finally, we summarized our proteome data on a model of dopaminergic transmission inspired from the KEGG database (<u>Mmu04728</u>) to represent proteins either enriched, retained or absent from our screen (Figure 3I Table S3 S5).

### Validation of 6 new proteins enriched at dopamine synapses

To further validate our proteomics screen, we monitored the fractionation of 6 candidates after DA-FASS using immunofluorescence. Copine7 is a C2 domain-containing, calcium-dependent, phospholipid-binding protein (Cpne7; 1.72-fold enrichment measured in MS/MS see Fig 3F and Table S3), displays a strong expression in dopaminergic cells of the midbrain, but also a significant expression in cholinergic interneurons of the striatum (CIN) and in potential cortico-striatal cells(Creutz *et al*, 1998; Savino *et al*, 1999) (Figure 4A and S3). We found Copine7 either colocalised or apposed to Th positive synaptosomes in 8% of labelled synaptosomes, a percentage that was maintained through DA-FASS purification (Figure 4B). Mint1/Apba1 (Mint1 for Munc18-1 interacting protein 1, also known as Amyloid Beta Precursor Protein Binding Family A Member 1; 1.57-fold enrichment measured in MS/MS see Fig 3F and Table S3) is a neuronal adapter protein that interacts with the Alzheimer’s disease amyloid precursor protein (APP) and plays a role at the synaptic active zone of neurotransmitter release (Miller *et al*, 2006; Südhof, 2012). Mint1/Apba1 was also shown to be involved in amphetamine-induced dopamine release (Mori *et al*, 2002). Mint1/Apba1 mRNA displays a strong expression in Th cells of the midbrain and a milder expression in CIN and potential cortical and thalamic afferent neurons (Figure 4A and S3). We found Mint1/Apba1 either colocalised or apposed with Th positive particles in 4% of all labelled synaptosomes, a percentage that was increased to 9% upon DA-FASS purification (Figure 4C). Cadps2 (Calcium-Dependent Activator Protein For Secretion 2; 1.62-fold enrichment measured in MS/MS see Fig 3F and Table S3) has been shown to play an important role in neurotransmitter secretion and monoamine loading in vesicles (Jockusch *et al*, 2007; Ratai *et al*, 2019). mRNA expression of Cadps2 is high in Th positive cells and significant in putative cortico-striatal cells (Figure 4A and S3)(Speidel *et al*, 2003). Indeed, we found Cadps2 both colocalized or apposed with Th signals in 13% of all labeled synaptosomes, a rate increased to 21% after sorting (Figure 4D). SynCAM 2 (SynCAM 2 for Synaptic cell adhesion molecule 2 also known as CADM2 for Cell adhesion molecule 2; 1.28-fold not significant enrichment ratio) is thought to mediate heterophilic trans-synaptic adhesion at excitatory synapses (Fogel *et al*, 2007; Thomas *et al*, 2008). While SynCAM 2 mRNA is highly expressed in all populations of neurons constituting the striatal neuropil, it is striking that SynCAM 2 expression is the highest in the brain in a subcluster of Th positive cells of the midbrain (Figure 4A, S3) (Saunders *et al*, 2018). Hence, SynCAM 2 represents an interesting candidate to promote synaptic adhesion at dopamine hub synapses. SynCAM 2 was seen mostly colocalized but also closely apposed with DAT signals in 28% of all labeled synaptosomes, a rate steeply increased to 75% after sorting (Figure 4E). Interestingly, SynCAM2 is associated to dopamine synaptosomes at a level comparable to Th (see Figure 2) but it is not a selective marker as it is expressed at many other synapses. Stx4 (Syntaxin 4; 3.36-fold enrichment measured in MS/MS see Fig 3F and Table S3) is a SNARE protein (soluble N-ethylmaleimide-sensitive factor attachment protein receptor) shown to mediate steps of membrane recycling at dendritic spines (Kennedy *et al*, 2010; Arendt *et al*, 2015). mRNA expression of Stx4 is moderate throughout afferent and efferent cells of the striatal neuropil (Figure 4A and S3). Stx4 signals were mostly apposed to Th signals in 7% of all labeled synaptosomes, a rate increased to 33% after sorting (Figure 4F). Finally, Mgll (Monoglyceride lipase; 1.93-fold enrichment measured in MS/MS see Fig 3F and Table S3) catalyzes the conversion of monoacylglycerides to free fatty acids and glycerol and is involved in the catabolism of the endocannabinoid 2-AG (2-arachidonoylglycerol) (Dinh *et al*, 2002). Mgll mRNA was detected at mild to high levels in most cell types afferent or efferent to the striatum, but the lowest expressers were the dopaminergic cells of the midbrain (Figure 4A and S3). Indeed, we found Mgll apposed to Th signals in 3% of all labelled synaptosome a percentage that increased to 9% upon sorting (Figure 4G).

**Figure 4.**
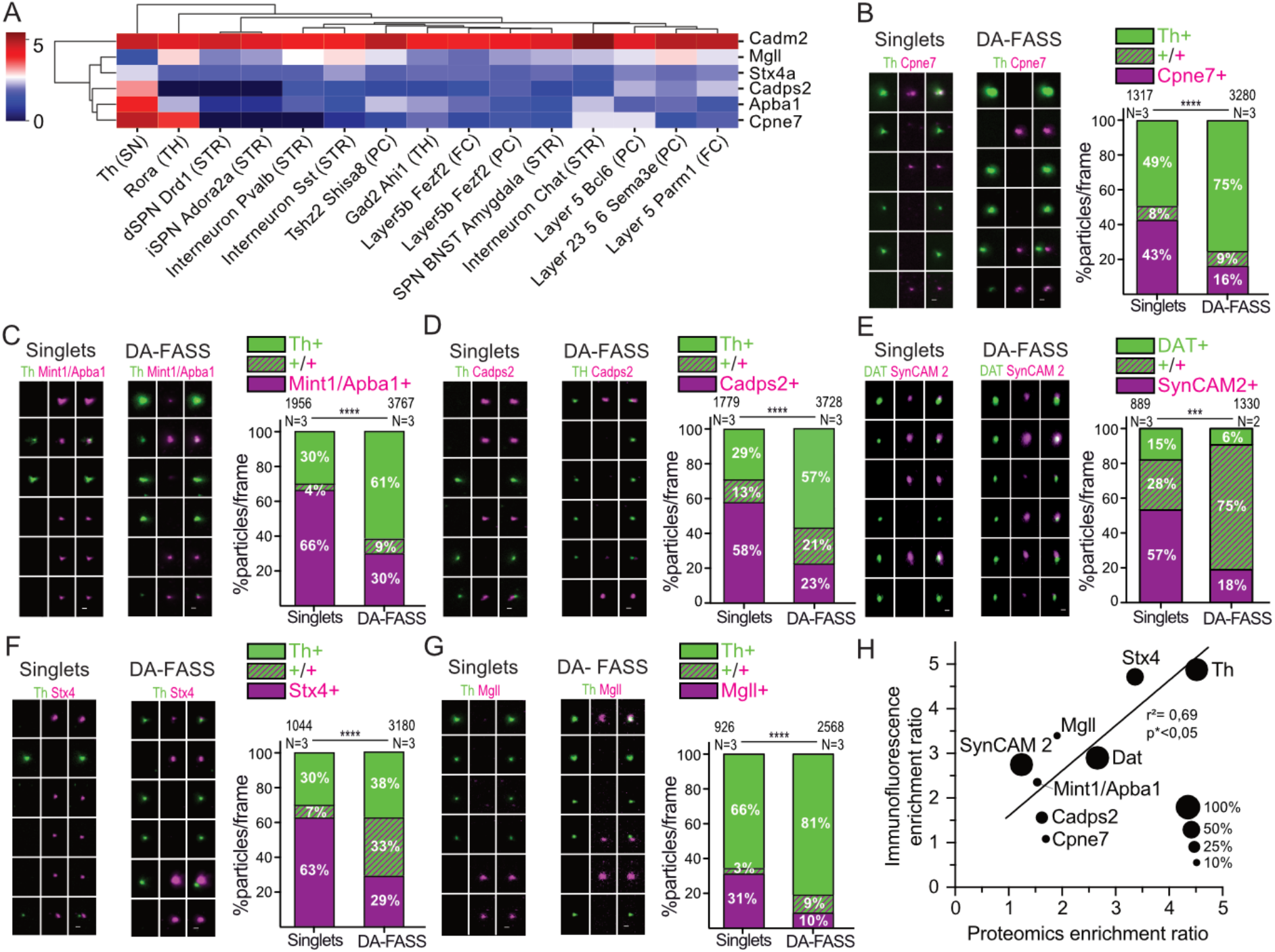
Validation of a selected set of DA-FASS enriched proteins with immunofluorescence. **(A)** Heatmap showing cell type specific mRNA abundance of the 6 enriched DA-FASS proteins selected for further experimental validation (detailed from figure 3H). **(B-G)** Epifluorescence images (left) of a representative sample of synaptosome populations (singlets left; DA-FASS right) labelled with anti-Th (green) and anti-Cpne7 (B), Mint1/Apba1 (C), Cadps2 (D), SynCAM 2 (E), Stx4 (F), Mgll (G) (magenta). (right) Analysis of staining showing particle proportions per frame. mean, interaction ****p < 0.001. Two-way MD ANOVA. Double positive particles population of most selected targets display above two folds enrichments (Mint1, C; Cadm2, E; Mgll, G) up to above four folds (Stx4, F) in the DA FASS sample. CPNE7, the least enriched protein in proteomics stay stable at around 9% of double positive particles with Th (C). Note that single positive particles are depleted upon DA FASS sorting for all selected proteins. Complete statistics are available in supplementary table 2. For all panels, scale bar = 1 μm. **(H)** Correlation between double positive immunofluorescence particle count and label free mass spectrometry based enrichment ratios (p<0.05, Pearson’s correlation coefficient r^2^=0.69). Dot sizes are scaled to the proportion of dopaminergic synaptosomes expressing each marker.

Altogether we could validate 6 new proteins from our screen for their selective association with dopaminergic synaptosomes. Interestingly, a comparison between MS/MS label free quantification and the immunofluorescence particle count association reveals a very good linear correlation of the results (Figure 4H).

### Proteins retained during DA-FASS delineate the association of dopaminergic varicosities in hub synapses

To investigate the identity of partners in the synaptic hubs we pursued the comparison of our screen with neurotransmission pathways reported in KEGG (Figure 3I). Exploring the pathway of SV and neurotransmitter cycling reveals a very high coverage of our proteome with proteins involved in neurotransmitter release, SV endocytosis and neurotransmitter uptake (Figure 5A gray boxed text, Table S3 S5). To complete this observation we probed for the phospho-proteins Synapsin 1&2 that are found at all presynapses (De Camilli *et al*, 1983) (abundance ratio 1.03 for both isoforms in our screen). EGFP+/synapsin+ particles representations rose from 8% to 45% upon DA-FASS while EGFP-/Synapsin+ synaptosomes were reduced from 84% to 35% after sort (Figure 5B). Synapsins were very frequently apposed to DAT-EGFP signals, an observation similar to the EM observation of multiple varicosities at synaptic hubs.

**Figure 5.**
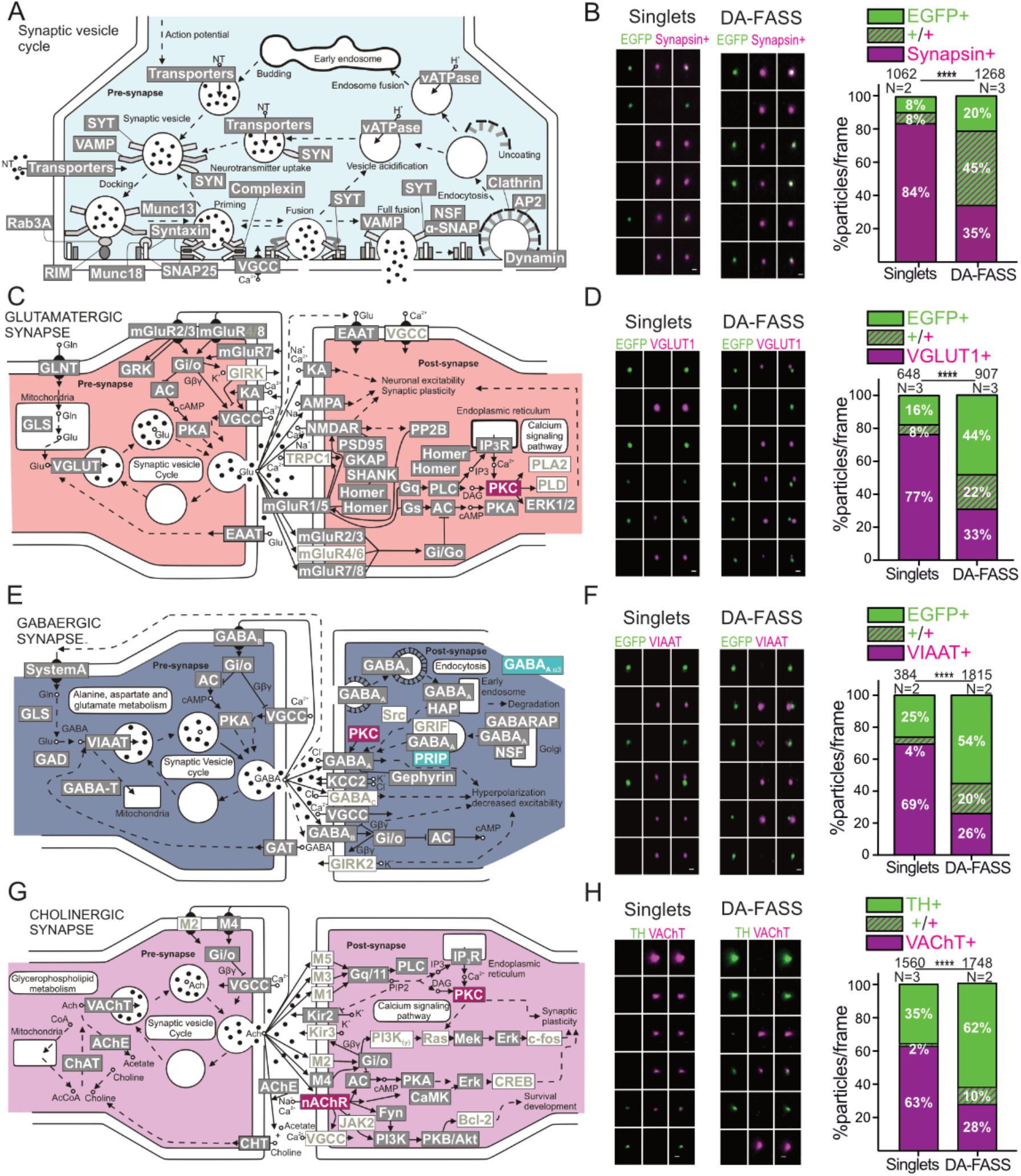
Proteomics and immunofluorescence of DA FASS sample reveals dopamine synapse association with other synaptic partners. **(A-C-E-G)** Schematic of the molecular organization of the synaptic vesicle cycle (A), glutamatergic (C), GABAergic (E) and cholinergic (G) synapses (Adapted from the database KEGG). Enriched proteins from our DA FASS sample are in red, depleted in cyan, retained in grey, and absent in white. Gene names for each protein can be found in supplementary table 4 (ST4). **(B-D-F-H)** Epifluorescence images (left) of a representative sample of synaptosome populations (singlets left; DA-FASS right) labelled with anti-EGFP or anti-Th (green) and anti-Synapsin (B), VGLUT1 (D), VIAAT (F) and VAChT (H) (magenta). (right) Quantification of stainings showing particle proportions per frame. mean, interaction ****p < 0.001. Two-way MD ANOVA. Note that for all vesicular synaptic markers, single positive particles are depleted upon DA FASS sorting while double positive particles are enriched. Complete statistics are available in supplementary table 2. For all panels, scale bar = 1 μm. See VGLUT2 immunofluorescence analysis in Figure S4.

We then explored the proteome related to excitatory synapses. Our coverage seems reliable because most categories of proteins are kept through DA-FASS (Figure 5C gray boxed text, Table S3 S5). We probed for the 2 vesicular glutamate transporters (VGLUT) to identify VGLUT1 expressing excitatory cortico-striatal inputs impinging on spines of the spiny projection neurons (SPNs) and VGLUT2 expressing thalamo-striatal inputs contacting SPNs (Herzog *et al*, 2001; Moss & Bolam, 2008). VGLUT1 varicosities were apposed to EGFP varicosities at hub synaptosomes. Through DA-FASS, EGFP-/VGLUT1+ synaptosomes were depleted more than 2-fold (from 77% to 33% sorted; Figure 5D). Yet, a third of dopaminergic EGFP+ synaptosomes were associated with a VGLUT1 pre-synapse (EGFP+/VGLUT1-44%, EGFP+/VGLUT1+ 22%; Figure 5D), and enriched almost 3-fold through DA-FASS (from 8% to 22%; Figure 5D). VGLUT2 signals followed the same trend with yet a lower percentage of association than the one reached with VGLUT1 (13% of DA positive synaptosomes; Figure S4). As a control we tested whether the reverse FASS experiment, sorting VGLUT1^venus^ cortico-striatal synaptosomes would co-purify VGLUT2-labelled terminals (see Figure S5AG). As expected, VGLUT2 synaptosomes were mostly segregated from VGLUT1^venus^ positive synaptosomes. Decisively, VGLUT2+/VGLUT1^venus^ + particles were not co-enriched through fluorescence sorting (9% in unsorted sample vs 6% in sorted synaptosomes; Figure S5FG upper right quadrants). This absence of segregation is consistent with the fact that these 2 markers were shown to contact distinct spines on SPNs (Moss & Bolam, 2008; Doig *et al*, 2010; Heck *et al*, 2015).

Our proteome also displays an abundant representation of markers of inhibitory synapses kept through DA-FASS enrichment (Figure 5C gray boxed text, Table S3 S5). Of note, 2 proteins of GABAergic synapses were seen depleted (Gabra3 and Prip; Figure 5E blue boxed text, Table S3 S5). We therefore probed for the vesicular inhibitory amino-acid transporter (VIAAT), that labels GABAergic terminals arising from all inhibitory neurons of the striatum (Sagné *et al*, 1997). VIAAT+/EGFP+ hub synaptosomes displayed 5-fold enrichment through DA-FASS (from 4% to 20%; Figure 5F), while the EGFP-/VIAAT+ population was depleted more than 2-fold (69% in unsorted vs 26% in sorted samples; Figure 5F). Hence, GABAergic synaptosomes were associated to 27% of the dopaminergic synaptosomes.

Finally striatal neuropils harbour a dense innervation by local cholinergic interneurons that function in tight interrelation with dopaminergic signals (Threlfell *et al*, 2012; Kitabatake *et al*, 2003). In accordance, our proteome also displays a significant fraction of cholinergic markers that are kept through DA-FASS (Figure 5G gray boxed text, Table S3 S5). The beta2 nicotinic receptor subunit (Chrnb2) is even significantly enriched (Figure 5G red boxed text, Table S3 S5). Indeed, it was shown to mediate cholinergic signalling onto dopaminergic varicosities (Wonnacott *et al*, 2000). To confirm a physical binding of dopaminergic varicosities with cholinergic ones, we probed for the Vesicular Acetyl Choline Transporter (VAChT). VAChT signals were occasionally seen apposed to Th positive dots with a 6-fold increase through DA-FASS enrichment (from 2% to 13%; Figure 5H). Through DA-FASS, Th-/VAChT+ synaptosomes were depleted nearly 2-Fold (from 63% to 35%; Figure 5H).

Hence our molecular data supports a frequent association of dopamine synapses with all the major synaptic partners operating in striatal neuropil supporting our earlier electron microscopy observations. To challenge the accuracy and specificity of our observations, we performed several controls. A test was applied to our images in order to established the random probability of co-sedimentation of separate particles at the same sites (see methods). Indeed, for all our datasets, random associations occurred on less than 2% of all events while we observed at least 11% for synaptic hub related associations in sorted samples (see Table 1). As a final control for the specificity of hub-synaptosome adhesion, we performed an additional VGLUT1^venus^ FASS experiment through which we selectively sorted aggregates and performed electron microscopy on our aggregate sorted sample (Figure S5 H-J). Upon reanalysis, sorted aggregates displayed a steep increase in the representation of small and large aggregates (Figure S5HI). Singlets were still strongly represented in the reanalysed sample as it is common to brake-down aggregates into singlets through the shearing forces applied in the nozzle of the FACS (Figure S5HI). Electron micrographs displayed many profiles of large particles (3-6µm in diameter) difficult to relate to identifiable features of the tissue and very different from the synaptosomes displayed in Figure 1 (Figure S5J).

**Table 1:** Observed versus simulated randomized associations of immunolabeled markers.

| Immuno-labelling | Observed associations (%) | simulated associations (%) |
| --- | --- | --- |
| EGFP + VGLUT1 | 22.4 | 0.94 |
| EGFP + VGLUT2 | 11.5 | 0.23 |
| EGFP + VIAAT | 20.2 | 0.67 |
| VGLUT1 + VGLUT2 | 5.45 | 1.98 |
| VGLUT1 + TH | 37.9 | 1.40 |
| TH + VACHT | 10,54 | 1,08 |

Altogether, we identified the association of dopaminergic and effector synapses in synaptic hub structures that specifically adhere together and may mediate the modulatory influence of dopamine over excitatory and inhibitory synaptic signalling. Cholinergic inputs from CINs may also take part to this association.

### Spatial organisation of dopaminergic synaptic hubs

Using a combination of distance measurements on the whole population of immunostained synaptosomes and STED microscopy we checked the relative position of each marker to dopaminergic varicosities (labelled against EGFP, Th or DAT). We first calculated the barycentre to barycentre distance between each markers and EGFP/TH/DAT immunofluorescence signals in DA-FASS samples. The accumulation of data over several hundred dots allowed for a fairly precise estimate of the distance between markers. TH is colocalized with EGFP and seen at an average distance equivalent to the resolution of our conventional epifluorescence setup (0.250µm), while the most distant marker, VGLUT1, is apposed on average at 0.639µm distance from EGFP+ barycentre (see Figure 6A-B). EGFP+/Synapsin+ synaptosomes represented 70% of all EGFP+ synaptosomes (Figure 5B). A careful scrutiny using STED microscopy shows that some EGFP+ synaptosomes contain both co-localized and apposed synapsin labels (Figure 6C). Apposed Synapsin puncta may represent glutamatergic, GABAergic, or cholinergic presynapses. VGLUT1, VGLUT2, and VIAAT signals display a rather distant apposition to EGFP+ varicosities (Figure 6D-F). SynCAM 2 is seen at 92% of DAT positive synaptosomes (Figure 4E). In STED, SynCAM 2 patches were visible tightly apposed to the DAT varicosities (Figure 6G).

**Figure 6.**
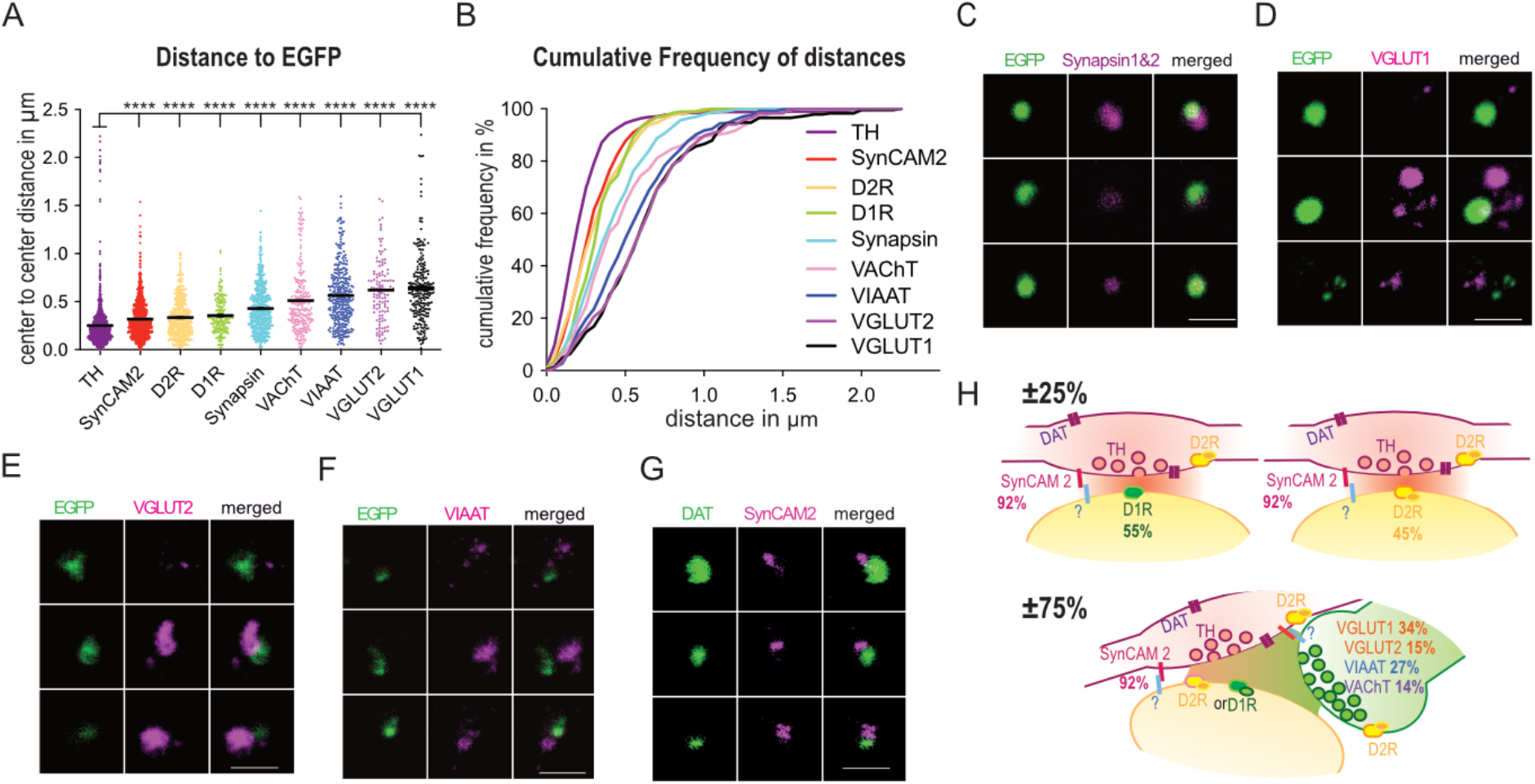
Modeling the spatial organization of dopaminergic synaptic-hubs. **(A)** Distance to EGFP barycenter for all stained proteins in increasing order of average distance (Th reference for VAChT, DAT reference for SynCAM2). Mean ± SEM, ****p<0,0001. Kruskal-Wallis. **(B)** Cumulative frequency distribution of distances to EGFP (percentage of all stained molecules) (Th reference for VAChT, DAT reference for SynCAM2). Note the proximity of SynCAM2 and the larger gap left by VAChT, VGLUT and VIAAT varicosities. **(C-G)** STED images of synaptosomes stained for EGFP (C-F) or DAT (G) (green) and Synapsin 1&2 (C), VGLUT1(D), VGLUT2 (E), VIAAT (F), SynCAM2 (G) (magenta). Scale bars 1 µm. **(H)** Synaptomic model of the dopamine synapse population in the striatum. Dopamine varicosities labelled by the DAT-cre strategy are apposed to a postsynaptic element with either D1R (55%) or D2R (45%). Present in nearly all boutons (92%) SynCAM 2 represents a good candidate for synaptic adhesion. Roughly, 75% of dopamine varicosities additionally form a synaptic hub with effector synapses (VGLUT1+ in 34%, VGLUT2+ in 13%, VIAAT+ in 27%). Additionally some association with cholinergic terminals is seen in 14% of the cases.

We thus propose a synaptomic model in which most dopaminergic varicosities adhere through SynCAM 2 to post-synaptic elements labelled by either D1 or D2 receptors. Beyond, a majority of dopaminergic synapses is also associated to effector synapses in synaptic hub structures clearly identified in electron and STED microscopy (Figure 6H).

### Dopaminergic innervation strengthens VGLUT1 excitatory cortico-striatal synapses

Finally, we wondered if the apposition of a dopaminergic innervation affect effector synapses. To that end we sorted striatal VGLUT1^venus^ synaptosomes with FASS (Figure 7A and Figure S5 A-E), stained them with Th to classify them into Th+ and Th-, and probed several markers of glutamatergic synapses. In this experiment, we confirm previous data (See Figure 5D) that VGLUT1+/Th+ particles are enriched with FASS and represent 40.8% of the VGLUT1+ population (Figure 7B). We further confirmed this using multiplexed capillary electrophoresis-based immunoblots with both VGLUT1^venus^ and Th detections in the same capillary. Th levels are maintained in FASS while VGLUT1^venus^ is enriched (Figure 7C). Hence, striatal VGLUT1^venus^ FASS sample is ideally suited to probe for a comparison between stand-alone VGLUT1 cortico-striatal synapses (Th-) and dopaminergic hub cortico-striatal synapses (Th+). VGLUT1^venus^ reports for the SV cluster and the loading of SV with glutamate(Herzog *et al*, 2011). We found a significantly higher VGLUT1^venus^ signal in Th+/VGLUT1+ compared to Th-/VGLUT1+ synaptosomes (143489 ± 2037 N=3 n=3609 for Th-; 182796 ± 4513 N=3 n=1206 for Th+; see Figure 7D). Bassoon is a scaffold protein of the active-zone of neurotransmitter release (Altrock *et al*, 2003). A recent report showed that a third of dopamine varicosities harbour a cluster of Bassoon (Liu *et al*, 2018). We could confirm that most VGLUT1+ synaptosomes display Bassoon signal close to the active-zone (93% association; Figure S6A), however, when looking at Th+/VGLUT1-elements only 19% contained a bassoon cluster (Figure S6A). When monitoring the intensity of bassoon at VGLUT1 synapses, we found significantly higher bassoon signal in Th+/VGLUT1+ compared to Th-/VGLUT1+ synaptosomes (138149 ± 1856 N=3 n=3609 for Th-; 205934 ± 7061 N=3 n=1206 for Th+ see Figure 7D and Figure S6A). To discriminate the origin of the bassoon signal, we performed STED imaging. We found some Th+ synaptosomes with a bassoon cluster but most of the Th+/VGLUT1+ synaptosomes do not display bassoon signal within Th varicosities (Figure 7E).

**Figure 7.**
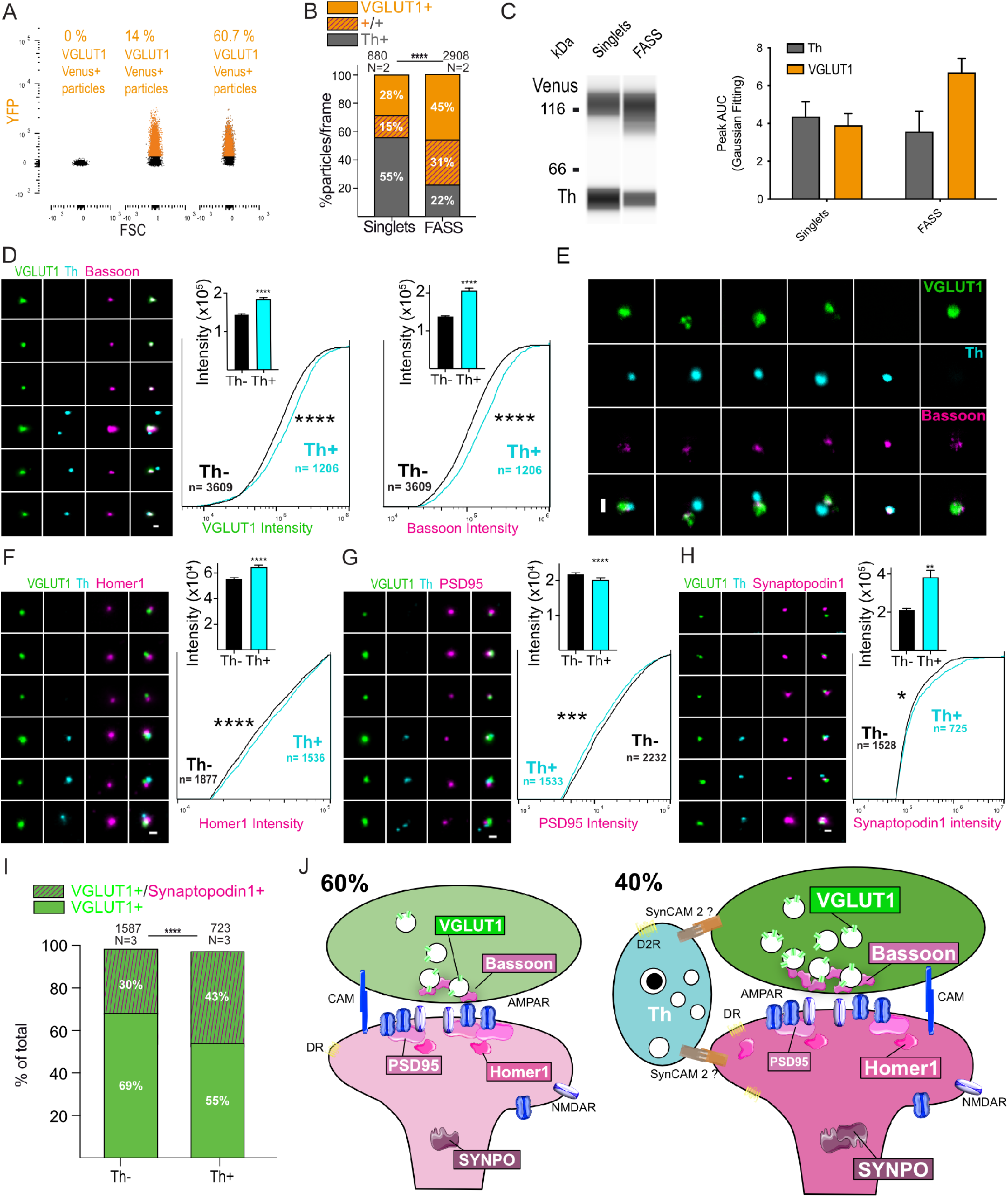
Hub synapse formation strengthen the molecular scaffolds of glutamatergic synapses. **(A)** VGLUT1^Venus^ FASS sorting from striatal explants. (left) Saline synaptosomes determine the level of autofluorescence. Gating was set to have 0% of particles within the VGLUT1^Venus+^ range in negative controls. (middle) Singlets VGLUT1^Venus+^ synaptosome before sorting show 14% of VGLUT1^Venus+^ particles. (right) FASS VGLUT1^Venus+^ synaptosomes re-analysed in the sorter consist of 60.7% of VGLUT1^Venus+^ particles. See Figure S5 for detailed gating strategy**. (B)** Analysis of staining of synaptosome populations (singlets; VGLUT1^Venus+^ FASS) labelled with anti-EGFP and anti-Th showing particle proportions per frame. mean, interaction ****p < 0.001. Two-way MD ANOVA. (**C)** (Right) Representative immunoblot bands of VGLUT1^Venus^ and Th. (Left) Averaged Th (grey) and VGLUT1^Venus^ (orange) peak AUC data, n=3 sorts, all data are mean ±SEM. Note that Th is maintained in the VGLUT1^Venus+^ FASS sample (C) and 40% of VGLUT1 synaptosomes are Th+ (B). **(D-F-G-H)** Representative epifluorescence images of VGLUT1^Venus+^ FASS synaptosomes immunolabelled for VGLUT1^Venus^ (green), Th (cyan), and respectively Bassoon (D), Homer1 (F), PSD95 (G) and synaptopodin1 (H) (magenta). Analysis of mean intensity as well as Cumulative Distribution Frequency of intensities between Th+ hub synapse and Th-VGLUT1 synapse populations are revealing a higher intensity of VGLUT1^Venus^ (D, left), Bassoon (D, right), Homer1 (F) Synaptopodin1 (H) and a slightly lower intensity of PSD95 in the Th+ population. Intensities are expressed as Mean ± SEM, ****p<0,0001, ** p<0,01. Mann Whitney. CDF *p<0,05; ***p<0,001 ****p<0,0001. Kolmogorov Smirnov. **(E)** STED images of VGLUT1^Venus^/TH/Bassoon synaptosomes. Note that in hubs, Bassoon signal is mostly found within VGLUT1^Venus^ signal when compared and not with Th. **(I)** Analysis of VGLUT1^Venus^ + and Synaptopodin1 + staining in presence or absence of Th showing particle proportions per frame. Mean, interaction ****p < 0.001. Two-way MD ANOVA. **(J)** Synaptomic model of VGLUT1 striatal synapses. 40% of VGLUT1 synapses are associated to a dopaminergic hub. In presence of Th positive varicosities, VGLUT1 synapses are strengthened as shown by an increased intensity of VGLUT1, Bassoon, Homer1 and Synaptopodin1. Some rearrangement may occur at the post-synapse with a slight decrease of PSD-95. For all panels scale bar = 1 μm.

Th+ varicosities do not seem to recruit more bassoon when they are taking part to a synaptic hub with VGLUT1 synapses. Hence it is reasonable to consider that most of the increase of bassoon signal at hub synapses is occurring at VGLUT1 terminal under the influence of dopaminergic innervation.

We then focused on post-synaptic proteins. Homer1 is a calcium binding scaffold protein important for metabotropic glutamate receptor signalling (Fagni *et al*, 2002). We found higher Homer1 signal in Th+/VGLUT1+ compared to Th-/VGLUT1+ synaptosomes (55356 ± 1052 N=3 n=1877 for Th-; 64756 ± 1385 N=3 n=1536 for Th+; see Figure 7F). In contrast, we found that the signal for PSD-95, a major post-synaptic density scaffold (Kim & Sheng, 2004), was slightly decreased in Th+/VGLUT1+ compared to Th-/VGLUT1+ synaptosomes (21758 ± 471,1 N=3 n=2232 for Th-; 20149 ± 580,9N=3 n=1533 for Th+; see Figure 7G). To further characterize the spine compartment, we then looked at synaptopodin (Synpo) a marker of the spine apparatus (Mundel *et al*, 1997). The spine apparatus and Synpo are found at a minority of spines in the forebrain and is thought to be recruited upon morphological potentiation event (Vlachos, 2012). When monitoring the intensity of signals, we found nearly twice more synpo at Th+/VGLUT1+ compared to Th-/VGLUT1+ synaptosomes (213786 ± 6158 N=3 n=1528 for Th-; 382663 ± 36567 N=3 n=725 for Th+; see Figure 7H). Surprisingly, we also observed that Synpo is relatively well maintained through FASS purification in the striatum while we had shown previously a strong depletion in forebrain samples (Biesemann *et al*, 2014; Hafner *et al*, 2019). In fact, Synpo is among the markers seen specifically enriched in the striatum in mass spectrometry compared to other brain regions (Sharma *et al*, 2015). In our LC-MS/MS screen, Synpo is unchanged after FASS (abundance-ratio of 1 with 17 unique peptides, Table S3), a trend we could confirm with immunofluorescence probing of synpo on DA-FASS samples (Figure S6B). In the triple staining experiment with VGLUT1^venus^ FASS, Synpo signals were apposed to VGLUT1 positive dots more frequently when Th positive varicosities were present (from 43% to 30%; Figure 7I).

Altogether our results show that a selective sets of markers of SV cluster, active-zone, post-synaptic density and spine apparatus at VGLUT1 synapses on SPNs display a strong increase upon innervation by dopaminergic varicosities.

## DISCUSSION

In order to unravel specific molecular and cellular features of modulatory neurotransmission, we targeted the dopaminergic projection from substantia nigra and ventral tegmental area to the striatum using FASS (Biesemann *et al*, 2014; Zhu *et al*, 2018). Specificity for dopaminergic synaptosomes was validated by the enrichment for presynaptic dopaminergic markers as well as the adhesion of dopaminergic varicosities to post-synaptic elements containing D1R or D2R. We could produce a proteome that quantifies 2653 proteins among which 57 are significantly enriched through DA-FASS purification. We validated 6 proteins identified in our screen (Cpne7, Mint1/Apba1, Cadps2, Cadm2/SynCAM 2, Stx4, Mgll). We show the association of dopaminergic synapses with glutamatergic, GABAergic and cholinergic synapses in structures identified in electron microscopy that we propose to name “dopaminergic hub synapses”. Finally, we observed that innervation of glutamatergic synapses with dopaminergic varicosities induces a molecular strengthening of the whole synapse.

### Cellular organisation of dopaminergic projections to the striatum

The nature of dopaminergic synaptic structures is the topic of a long-standing debate. Previous anatomical investigations in the field identified that the distribution of dopamine varicosities in the neuropil is biased toward a proximity to effector synapses but only a minority makes synapses with a target structure in the striatum (Kreitzer, 2009; Descarries *et al*, 1996; Moss & Bolam, 2008). However, other authors reported the frequent occurrence of symmetrical synaptic contacts of dopaminergic thin axonal portions with SPNs spines or dendritic shafts (Descarries *et al*, 1996; Freund *et al*, 1984; Moss & Bolam, 2008; Groves *et al*, 1994; Bamford *et al*, 2004; Gaugler *et al*, 2012). Our current dataset strongly advocates for a specific and frequent adhesion of dopaminergic axonal varicosities of rather small diameter with target structures (Figures 1H-L and 2). Indeed, around 55% of our EGFP+ varicosities displayed apposed D1R, while more that 80% displayed D2R labelling (Figure 2). This is in accordance with SPNs being the main target of dopamine terminals in the striatum, with roughly half of the SPNs expressing D1 receptors, while the D2 receptor is expressed by the other half (Bertran-Gonzalez *et al*, 2010; Calabresi *et al*, 2014) as well as by dopaminergic and effector presynapses (Sesack *et al*, 1994; Delle Donne *et al*, 1997; Hersch *et al*, 1995).

Moreover, our data reveals that adhesion at the dopaminergic varicosity extends to synaptic hubs with effector synapses. We found that around a third of dopaminergic varicosities make hub synapses with cortico-striatal VGLUT1 synapses, around 15% associate with thalamo-striatal VGLUT2 synapses, and more than a quarter was associated to VIAAT inhibitory synapses (Figures 5 and 6H). Additionally, around 14 % are also contacted by cholinergic inputs. Conversely, nearly half of VGLUT1 striatal synaptosomes were observed in hub synapses (Figure 7B). Providing that little overlap exists between those hub associations, more than 75% of dopaminergic varicosities may adhere to hub synapses. Indeed, VGLUT1 and VGLUT2 synaptosomes displayed little to no association when probed from a striatal sorting from VGLUT1^venus^ mice (Figure S5FG). According to the literature, cholinergic inputs to these hubs may target dopaminergic varicosities(Threlfell *et al*, 2012; Wonnacott *et al*, 2000), further investigations will be required to determine this in detail. As we could show that most of the soluble content of dopaminergic axons is engulfed in synaptosomes of our preparation (Figure 1B-D), we feel confident to state that most dopaminergic varicosities engage into hub synapses, while asynaptic varicosities may represent a minority in the tissue. Dopaminergic varicosities are clear to electrons and less populated with SVs (Moss & Bolam, 2008; Freund *et al*, 1984; Gaugler *et al*, 2012). Synaptic hubs observed here associate electron dense terminals strongly populated with SVs, to clear varicosities much less populated with SVs (Figure 1FH). The occurrence of synaptic hubs may explain previous observations that striatal dopaminergic synaptosomes sediment faster than other synaptosomes in a linear sucrose gradient (Van der Krogt *et al*, 1983). Further investigations will be necessary to unravel whether synaptic hub formation is a structural invariant common to all sub-divisions of the striatum, and whether the proportion of hub synapses of different kinds can vary depending on subregions and/or physiological states.

### A proteome of dopaminergic synapses in the striatum

Our first molecular characterization of FASS dopaminergic synaptosomes identifies 2653 proteins quantified between unsorted synaptosomes and DA-FASS samples. The structure of the data shows that 63 proteins are significantly depleted during the DA-FASS procedure while 57 proteins are seen strongly enriched. Hence, most proteins are kept in the process. This may be attributed to the existence of hub synapses with most other neuronal partners in the striatum on the one hand, and to the relative low purity of our DA-FASS samples until now on the other hand (more than 40% of contaminants are left after sorting according to reanalysis, Figure 1 and 3). Yet, the enrichment factor for dopaminergic synaptosomes is quite high (between 5- and 10-fold) which provides the opportunity to detect proteins selectively targeted to hub synapses. With the validation of 6 targets using immunofluorescence assay we show that our screen quality is high even for proteins with a low enrichment factor like copine 7 (Figure 4). Some proteins may populate presynapses of the hub partners (Cpne7, Mint1/Apba1, Cadps2, Cadm2/SynCAM 2, Mgll) while Stx4 may also be post-synaptic proteins based on the expression profile of the mRNA (Saunders *et al*, 2018) and previous publications (Mori *et al*, 2002; Speidel *et al*, 2003; Thomas *et al*, 2008; Dinh *et al*, 2002; Kennedy *et al*, 2010). Cpne7, Mint1/Apba1, Cadps2 and Stx4 point to specific membrane trafficking features at dopamine hub synapses (Creutz *et al*, 1998; Mori *et al*, 2002; Ratai *et al*, 2019; Kennedy *et al*, 2010). The cross analysis of our screen with single cell RNA sequencing data allowed us to spot the synaptic adhesion protein SynCAM 2 (Cadm2; Figure 3 and Figure S3) as a potential important player of adhesion at dopaminergic varicosities (Biederer *et al*, 2002; Biederer, 2006; Fogel *et al*, 2007). Our immunofluorescence data confirms the strong expression of SynCAM 2 at dopaminergic varicosities (Figure 4E, 6ABG). SynCAM 2 was also reported to label axons (Pellissier *et al*, 2007). We therefore propose that SynCAM 2 is part of an axonal adhesion complex responsible for the formation of dopaminergic synapses with SPNs and hub synapses with effector moieties. SynCAM 2 is thought to engage in heterophilic interactions with SynCAM 1 or 4 (Thomas *et al*, 2008; Fogel *et al*, 2007). Interestingly, SynCAM 1 is a player of cocaine-induced synaptic plasticity in the striatum (Giza *et al*, 2013) and SynCAM 2 regulates food intake and energy balance (Yan *et al*, 2018), two phenomena directly related to the dopaminergic system (Tellez *et al*, 2013, 2016). Besides, SynCAM 1 is thought to be preferentially acting at the post-synapses to induce presynaptic adhesion (Czöndör *et al*, 2013; Fowler *et al*, 2017). Hence, SynCAM 1 and 2 are strong candidates to mediate adhesion through heterophilic interaction at dopamine synapses, in addition to roles at other types of synapses. A previous contribution suggested that adhesion at dopaminergic synapses occurs through neuroligin 2 (Uchigashima *et al*, 2016). Our mass spectrometry data identified all 4 neuroligins and all 3 neurexins without specific enrichment through DA-FASS. Additionally, we did not find a selective enrichment for neuroligin 2 mRNA in SPNs that may justify to evaluate this protein further compared to SynCAM 2 (Saunders *et al*, 2018). The action of neuroligin 2 is preeminent at inhibitory synapses (Poulopoulos *et al*, 2009; Varoqueaux *et al*, 2004). One possible explanation could be that in the context of synaptic hubs, neuroligin 2 plays a role in the association with inhibitory synapses. Also, some work suggests a direct inhibitory function of dopamine projection on SPNs with mechanisms that still largely escape our understanding (Tritsch *et al*, 2012; Chantranupong *et al*, 2020). Nlgn2 may have a function related to this inhibitory phenotype yet this seems to go without a significant enrichment of Nlgn2 with dopaminergic synaptosome purification. Finally, a combinatorial of synaptic adhesion molecules is certainly involved and further investigations will be important to clarify the complete machinery responsible for dopaminergic hub synapse formation and maintenance (Südhof, 2021).

### Dopaminergic input to cortico-striatal synapses induces a molecular differentiation

Beyond showing the existence of dopamine hub synapses, we identified that the binding of Th varicosities to cortico-striatal synapses results in an increase in VGLUT1, Bassoon, Homer1 and Synaptopodin and a modest decrease in PSD95. To our knowledge this is the first time a physical interaction of dopamine axons with their target is shown to induce molecular differentiation. We found that nearly all dopaminergic terminals seem to adhere to a post-synaptic element populated with cognate receptors. Yet, around 2/3 of the dopaminergic varicosities are thought to be “silent” at a given time in the striatum and do not contain the full active zone molecular complement (Pereira *et al*, 2016; Liu *et al*, 2018). In accordance, we found that less than 20% of Th varicosities contain Bassoon in our samples regardless of their involvement in hub synapses. The higher frequency of bassoon in previous studies may have been biased by the crowding of excitatory and inhibitory synapses around the dopaminergic varicosities observed (Liu *et al*, 2018). Our FASS isolation approach reduces significantly the density of material in direct proximity to the observed synaptosomes. It thus seems like the formation of a synaptic contact at dopaminergic varicosity is not sufficient to establish a functional release site but rather additional plasticity may be required to engage silent dopaminergic synapses into dopamine transmission. Conversely, it remains to be established whether the molecular potentiation we describe results from a structural effect related to the binding of a Th+ varicosity or to local dopamine release (Yagishita, Science, 2014). Finally, it will be important to establish the function of newly identified proteins such a Stx4 or Cadm2/SynCAM 2 in the differentiation process, and the ultrastructural correlate of the observed molecular plasticity.

The discovery of molecular differentiation at synaptic hubs linking dopaminergic and effector synapses provides a unique ex-vivo paradigm to study the complex interactions of receptors – through signalling crosstalk or heteromeric interactions – identified in the past decades (Liu *et al*, 2000; Cepeda & Levine, 2012; Cahill *et al*, 2014; Ladépêche *et al*, 2013). As previously published, we found that many D1R or D2R labelled particles were extra-synaptic(Caillé *et al*, 1996; Sesack *et al*, 1994). Yet, most EGFP labelled synaptosomes displayed an apposed D1R or D2R. Therefore, the question of the co-recruitment of glutamate or GABA receptors with dopamine receptors at synaptic hubs is raised and the plasticity of this recruitment upon reward-based learning and in pathologic processes will have to be established.

Beyond, D1R and D2R interactions with effector ionotropic receptors, Adenosine, cannabinoid, metabotropic glutamate receptors and muscarinic receptors are also important players of the striatal integration of cortical and thalamic inputs. The increase in homer1 suggests a potential involvement of metabotropic glutamate receptors in the differentiation process (Fagni *et al*, 2002). Also, both homer1 and synaptopodin were shown to be involved in calcium signalling regulation, a point of interest for future investigations (Fagni *et al*, 2002; Vlachos, 2012). Downstream targets of signalling such as ionic channels may also take part to the critical scaffolds at play (Surmeier *et al*, 2007; Calabresi *et al*, 2014). Thus, DA-FASS synaptosomes will be a powerful tool to identify key molecular signalling complexes for dopamine action on striatal networks.

Altogether, our work paves the way for a better understanding of dopaminergic synaptic transmission in physiology and pathology (Blumenstock *et al*, 2019). Future developments will allow a more thorough multi-omics (Poulopoulos *et al*, 2019) as proposed recently with other techniques (Hobson *et al*, 2021). More generally, results from our study and the work of Apóstolo and colleagues (Apóstolo *et al*, 2020) on mossy fiber terminals of the hippocampus show that FASS synaptomics is a powerful workflow for the exploration of projection-specific synaptomes (Zhu *et al*, 2018; Grant, 2019).

## METHOD

### Animals

A transgenic mouse line expressing *cre* recombinase under the control of the dopamine transporter (DAT) was used (Turiault *et al*, 2007). Mice were maintained in C57BL/6N background and housed in 12/12 LD with ad libitum feeding. Every effort was made to minimize the number of animals used and their suffering. The experimental design and all procedures were in accordance with the European guide for the care and use of laboratory animals and approved by the ethics committee of Bordeaux University (CE50) and the French Ministry of Research under the APAFIS n° 8944 and #21132.

### AAV Vector and stereotaxic injection

Stereotaxic injections were performed in heterozygous DAT-*cre*^+^ and wild-type (WT) mice of either sex at 8 to 11 weeks of age (Cetin *et al*, 2006). An Adeno-Associated Virus (AAV) containing an inverted sequence of EGFP (AAV1 pCAG-FLEX-EGFP-WPRE, University of Pensylvania) (Oh *et al*, 2014) or mNeongreen (AAV1 pCAG-FLEX-mNeongreen-WPRE) (Shaner *et al*, 2013) coding gene flanked by loxP-sites was injected into DAT-*cre*^+^ mice (Figure 1 Panel 1). Saline injected littermates were used as auto-fluorescence controls. The stereotaxic injections were performed in Isoflurane-anesthetized mice using a 10 μl NanoFil syringe and a 35 G beveled NanoFil needle (World Precision Instruments). Injection coordinates for the Substantia Nigra pars compacta (SNc) were anterior/posterior (A/P) - 3.6 mm, lateral (L) +/− 1.3mm, dorsal/ventral (D/V) - 4.2mm. Injection coordinates for the Ventral Tegmental (VTA) were with a 12° angle A/P - 2.9mm, L +/- 1.6mm; D/V - 4.6mm. A/P and L coordinates are given with respect to the *bregma*, whereas D/V coordinates are given with respect to the brain surface (Figure 1 Panel 1). The animals were euthanized after 28 days at the maximal viral EGFP expression. For fluorescence activated synaptosome sorting (FASS) experiments, three to six DAT-*cre*^+^ mice and one WT mouse were used.

### Subcellular fractionation of synaptosomes

The preparation of sucrose synaptosomes was adapted from a previously published protocol (De-Smedt-Peyrusse *et al*, 2018). Briefly, animals were euthanized by cervical dislocation, decapitated and the head was immersed in liquid nitrogen for a few seconds. The striatum of WT and bright fluorescent parts of the striatum of DAT-*cre*^+^ mice were subsequently dissected under an epi-fluorescence stereomicroscope (Leica Microsystems, Germany, Figure 1 Panel 2). Non-fluorescent control striata were dissected following anatomical borders. Samples were then homogenized in 1ml of ice-cold Isosmolar buffer (0.32M sucrose, 4mM HEPES pH7.4, protease inhibitor cocktail Set 3 EDTA-free (EMD Millipore Corp.)) using a 2ml-glass-Teflon homogenizer with 12 strokes at 900 rpm. The homogenizer was rinsed with 250μL of isosmolar buffer and 3 manual strokes and then, the pestle was rinsed with additional 250μl of isosmolar buffer. The final 1.5ml of homogenate (H) was centrifuged at 1000xg for 5min at 4°C in a benchtop microcentrifuge. The supernatant (S1) was separated from the pellet (P1) and centrifuged at 12,600xg for 8min at 4°C. The supernatant (S2) was aliquoted and the synaptosomes-enriched pellet (P2) was resuspended in 350 µL of isosmolar buffer and layered on a two-step ficoll density gradient (5mL of 13% Ficoll, 0.32M sucrose, 4mM HEPES and 5mL of 7,5% Ficoll, 0.32M sucrose, 4mM HEPES). The gradient was centrifuged at 16,000 rpm for 1 hour and 10 min at 4°C (Thermo Sorvall WX Ultra 90 with a TH 641 rotor). The synaptosome fraction (Syn) was recovered at the 7.5 and 13% ficoll interface using a 0.5ml syringe. For complete subcellular fractionation 200 *μ* L of the P2 fraction was transferred to a 10 cm^3^ ice-cold glass/Teflon potter and quickly homogenized at full speed in 1.8 mL ultrapure water to create an osmotic shock. For synaptic vesicle fractionation the lysate was centrifuged at 25,000 × *g* for 7 min at 4 °C. The supernatant lysate (LS1) was extracted, the pellet (LP1) was resuspended in 200 µL of isosmolar buffer and part of it aliquoted. Remaining of LS1was centrifuged at 200,000 × *g* for 120 min at 4 °C. The whole supernatant LS2 was collected, concentrated to 50 µL and aliquoted. The pellet containing crude synaptic vesicles (LP2) was resuspended in 50 μL of ice-cold isosmolar buffer and aliquoted. LP1 fraction was centrifuged on the same discontinuous ficoll gradient (7.5 – 13%) at 60,000 × *g* for 33 min at 4 °C and the fraction at the interface of the two gradients containing synaptic plasma membranes (SPM) was collected and aliquoted.

### Fluorescence Activated Synaptosome Sorting (FASS) workflow

After collection, sucrose/ficoll synaptosomes were stored on ice and sequentially diluted in ice-cold PBS with protease inhibitor as described above, and the lipophilic dye FM4-64 dye was added at 1μg/ml to the solution to red label all membrane particles (Figure 1A). The FACSAria-II (BD Biosciences) was operated with the following settings: 70μm Nozzle, sample shaking 300rpm at 4°C, FSC neutral density (ND) filter 0.5, 488nm laser on, area scaling 1.18, window extension 0.5, sort precision 0-16-0, FSC (340 V), SSC (488/10nm, 365V), FITC (EGFP/mNeongreen) (530/30nm, 700V), PerCP (FM4-64) (675/20 nm, 700 V). Thresholding on FM4-64 was set with a detection threshold at 800. Samples were analyzed and sorted at rates of 18,000-23,000 events/s and flow rate of 3. Unsorted synaptosomes (“singlet” gate) and FASS synaptosomes (“EGFP+” or “mNeongreen+” or “VENUS+” sub-gate of the “singlet” gate) were collected sequentially (Figures 1, 7, S1 and S5). After sorting, samples were either centrifuged on gelatinized coverslips of 12mm diameter (5×10^5^ synaptosomes per coverslip at 6,800xg for 34min at 4°C Beckman J-26XP with a JS 5.3 rotor), or filtered on 0.1µm Durapore hydrophilic PVDF membranes (Merck-Milipore). Coverslips were then further treated and analyzed either for immunofluorescence imaging or for electron microscopy while filtered samples underwent, WES or mass spectrometry analysis (Figure 1A). Sorts’s statistical analysis was performed using two-way mixed design (MD) ANOVA (Figure 1F and S5E).

### Simple Western™ immunoblot

Detection proteins of interest were determined using WES (Simple Western™) by Protein Simple©. This system uses automated capillary electrophoresis based immunoblot to separate, identify and quantify a protein of interest. Reagents (Dithiothreitol, DTT; Fluorescent 5X Master Mix, Biotinylated Ladder) were prepared according to the manufacturer’s protocol. Samples were diluted with 0.1X Sample Buffer and mixed with 5X Master Mix (4 to 1) to obtain 50 ng/mL and finally denatured 5 min at 70°C. Primary antibodies were diluted to their tested optimal concentration and Luminol-Peroxide (1 to 1) mix was prepared. The plate was filled following the protocol scheme (5 µL of Biotinylated Ladder, 5 µL of Samples, 10 µL of Wes Antibody Diluent, 10 µL of Primary Antibody, 10 µL of Streptavidin-HRP, 10 µL of Secondary Antibody and 15 µL of Luminol-Peroxide Mix). Simple Western™ standard immunodetection protocol was run (separation matrix loading: 200 sec, stacking matrix loading: 15 sec, sample loading: 9 sec, separation: 25 min at 375 volts, antibody diluent: 5 min, primary antibody: 30 min, secondary antibody: 30 min, detection: high dynamic range). Capillary chemiluminescent images captured through a charge-coupled device camera were analyzed by the manufacturers Compass software. Briefly, the protein peaks area under the curve (AUC) were fitted using a Gaussian distribution. The fitted protein AUC is expressed either as a ratio to the fitted AUC H fraction for each WES (Figure 1) or as fitted protein AUC (Figure 7). WES’ data statistical analysis was performed using Two-way RM ANOVA (Table 2, Figure 1D and 7C).

### Immunofluorescence

Synaptosomes on coverslips were fixed (4% paraformaldehyde, 4% sucrose, PBS) for 10min at room temperature, washed three times with PBS for 5min and then stored at 4°C until use. Synapstosomes were blocked and permeabilized with PGT buffer (PBS, 2g/L gelatin, 0.25% Triton X-100 and when needed 5% normal goat serum) and subsequently incubated with primary antibodies in PGT buffer (1h at room temperature), washed 3 times with PGT and incubated with secondary antibodies in PGT (1 hour at room temperature). Three final washes with PGT buffer were performed prior to a washing step in PBS and a final rinse in ultrapure water. Coverslips were mounted on glass slides with Fluoromount-G mounting solution (Sigma) and stored at 4°C until observation.

### Antibodies

The following primary antibodies were used for immunofluorescence and WES. Anti-SynCAM 2 monoclonal antibody raised in rat against an epitope in the extracellular domain (provided by Thomas Biederer). Anti-D1 receptor, goat polyclonal antibody (Frontier Institute Cat# D1R-Go-Af1000, RRID: AB_2571594). Guinea pig polyclonal antibody to: VGLUT2 (Millipore Cat# AB2251, RRID:AB_1587626); VGLUT1 (Millipore Cat# AB5905, RRID:AB_2301751); MGL (Frontier Institute Cat# MGL-GP, RRID:AB_2716807); Homer 1 (Synaptic Systems Cat# 160 004, RRID:AB_10549720). Mouse monoclonal antibody to GFP (Roche Cat# 11814460001, RRID:AB_390913); Gephyrin (Synaptic Systems Cat# 147 111, RRID: AB_887719); Munc-18 (BD Biosciences Cat# 610336, RRID:AB_397726); Synaptophysin 1 (Synaptic Systems Cat# 101 011C3, RRID:AB_887822); Tyrosine Hydroxylase (Millipore Cat# MAB318, RRID:AB_2201528), PSD-95 (Abcam Cat# ab2723, RRID:AB_303248). Rabbit polyclonal antibodies to: GluA1 (Millipore Cat# AB1504, RRID:AB_2113602); Tyrosine Hydroxylase (Synaptic Systems Cat# 213 102, RRID:AB_2619896); D2 dopamine receptor (Millipore Cat# ABN462, RRID:AB_2094980); Synapsin 1/2 (Synaptic Systems Cat# 106 002, RRID:AB_887804); VGLUT2 (Cat# VGLUT2, RRID:AB_2315563)(Herzog *et al*, 2001); Sybnaptopodin (Synaptic Systems Cat# 163 002, RRID:AB_887825); VAChT (Synaptic Systems Cat# 139 103, RRID:AB_887864); VIAAT/VGATs (Synaptic Systems Cat# 131 002, RRID:AB_887871); DAT (Millipore Cat# AB2231, RRID: AB 1586991); EAAT1/GLAST (Cat# Ab#314, RRID:AB_2314561 a kind gift by Niels Christian Danbolt, University of Oslo)(Holmseth *et al*, 2009); GFP (Abcam Cat# ab290, RRID:AB_303395); Cpne7 (OriGene Cat# TA334534); Mint 1 (Synaptic Systems Cat# 144 103, RRID:AB_10635158); Cadps2 (Synaptic Systems Cat# 262 103, RRID:AB_2619980); Syntaxin 4 (Synaptic Systems Cat# 110 042, RRID:AB_887853);. Anti-Tyrosine hydroxylase, chicken antibody (Millipore Cat# AB9702, RRID:AB_570923).

### Proteomics

#### Sample preparation and protein digestion

Triplicates of 35*10^7^ FASS synaptosomes were accumulated for proteomics analysis and were compared to triplicates of 35*10^7^ singlets particles. Both samples were treated in parallel at all steps. Protein samples were solubilized in Laemmlli buffer. A small part of each triplicate was analysed by silver-staining using SilverXpress^R^ staining kit (Invitrogen, Cat#LC6100). Protein content was normalized across triplicates to 140 ng (lowest triplicate protein amount) and run onto SDS-PAGE (Sodium Dodecyl Sulfate-Poly Acrilamide Gel Ellectrophoresis) for a short separation. After colloidal blue staining, each lane was cut in 2 bands which were subsequently cut in 1 mm x 1 mm gel pieces. Gel pieces were destained in 25 mM ammonium bicarbonate 50% Acetonitrile (ACN), rinsed twice in ultrapure water and shrunk in ACN for 10 min. After ACN removal, gel pieces were dried at room temperature, covered with the trypsin solution (10 ng/µl in 50 mM NH_4_HCO_3_), rehydrated at 4 °C for 10 min, and finally incubated overnight at 37 °C. Spots were then incubated for 15 min in 50 mM NH_4_HCO_3_ at room temperature with rotary shaking. The supernatant was collected, and an H_2_O/ACN/HCOOH (47.5:47.5:5) extraction solution was added onto gel slices for 15 min. The extraction step was repeated twice. Supernatants were pooled and dried in a vacuum centrifuge. Digests were finally solubilized in 0.1% HCOOH.

#### nLC-MS/MS analysis and Label-Free Quantitative Data Analysis

Peptide mixture was analyzed on a Ultimate 3000 nanoLC system (Dionex, Amsterdam, The Netherlands) coupled to a Electrospray Orbitrap Fusion™ Lumos™ Tribrid™ Mass Spectrometer (Thermo Fisher Scientific, San Jose, CA). Ten microliters of peptide digests were loaded onto a 300-µm-inner diameter x 5-mm C_18_ PepMap^TM^ trap column (LC Packings) at a flow rate of 10 µL/min. The peptides were eluted from the trap column onto an analytical 75-mm id x 50-cm C18 Pep-Map column (LC Packings) with a 4–40% linear gradient of solvent B in 105 min (solvent A was 0.1% formic acid and solvent B was 0.1% formic acid in 80% ACN). The separation flow rate was set at 300 nL/min. The mass spectrometer operated in positive ion mode at a 1.8-kV needle voltage. Data were acquired using Xcalibur 4.3 software in a data-dependent mode. MS scans (*m/z* 375-1500) were recorded in the Orbitrap at a resolution of R = 120 000 (@ m/z 200) and an AGC target of 4 x 10^5^ ions collected within 50 ms. Dynamic exclusion was set to 60 s and top speed fragmentation in HCD mode was performed over a 3 s cycle. MS/MS scans were collected in the Orbitrap with a resolution of 30 000 and a maximum fill time of 54 ms. Only +2 to +7 charged ions were selected for fragmentation. Other settings were as follows: no sheath nor auxiliary gas flow, heated capillary temperature, 275 °C; normalized HCD collision energy of 30%, isolation width of 1.6 m/z, AGC target of 5 x 10^4^ and normalized AGC target od 100%. Advanced Peak Detection was activated. Monoisotopic precursor selection (MIPS) was set to Peptide and an intensity threshold was set to 2.5 x 10^4^.

#### Database search and results processing

Data were searched by SEQUEST through Proteome Discoverer 2.5 (Thermo Fisher Scientific Inc.) against the *Mus musculus* SwissProt protein database (v2021-02-04; 17,050 entries) added with the green fluorsecent reporter (mNeonGreen). Spectra from peptides higher than 5000 Daltons (Da) or lower than 350 Da were rejected. Precursor Detector node was included. Search parameters were as follows: mass accuracy of the monoisotopic peptide precursor and peptide fragments was set to 10 ppm and 0.02 Da respectively. Only b- and y-ions were considered for mass calculation. Oxidation of methionines (+16 Da), phosphorylation of serines, threonines and tyrosines (+79), methionine loss (−131 Da), methionine loss with acetylation (−89 Da) and protein N-terminal acetylation (+42Da) were considered as variable modifications while carbamidomethylation of cysteines (+57 Da) was considered as fixed modification. Two missed trypsin cleavages were allowed. Peptide validation was performed using Percolator algorithm (Käll *et al*, 2007) and only “high confidence” peptides were retained corresponding to a 1% False Positive Rate at peptide level. Peaks were detected and integrated using the Minora algorithm embedded in Proteome Discoverer. Proteins were quantified based on unique and razor peptides intensities. Normalization was performed based on total protein amount. Protein ratio were calculated as the median of all possible pairwise peptide ratios. A t-test was calculated based on background population of peptides or proteins. Quantitative data were considered for proteins quantified by a minimum of two peptides. Proteins with an abundance ratio above 1.5 were considered enriched and depleted below a ratio of 0.75 providing that data displayed a statistical p-value lower than 0.05.

The mass spectrometry proteomics data have been deposited to the ProteomeXchange Consortium via the PRIDE partner repository with the dataset identifier PXD027534 (Perez-Riverol *et al*, 2019)).

#### Meta-Analysis with former databases

Meta-analysis was carried out using databases from the mouse brain proteome (Sharma *et al*, 2015), SynGO (Koopmans *et al*, 2019), DropViz (Saunders *et al*, 2018). Volcano plots and heatmaps were created using python based bioinfokit (Bedre, 2021).

### Epifluorescence Microscopy and Image processing

Immuno-stained synaptosomes were imaged using either a Nikon Eclipse NiU (with a 40x/NA 0.75 dry objective equipped with a sCMOS ANDOR Zyla 5.5 sCMOS camera), a Leica DMI8 epifluorescence microscope (with a 63x/NA 1.4 oil immersion objective equipped with a sCMOS Hamamatsu FLASH 4.0v2 camera) or a Leica DM5000 epifluorescence microscope (with a 40x/NA 1.25 immersion objective equipped with a sCMOS Hamamatsu FLASH 4.0 camera; Figure 1). Ten to twenty frames were chosen randomly on each coverslip and imaged.

Correlation of synaptosomes’ labelling has been automated by generating a home-made macro-command, using the ImageJ software (Rasband, 1997) (18.02.19.Quantification.de.colocalisation.sur.synaptosomes.Herzog.Etienne.v5.ijm). The workflow is composed of three steps. First, the images are pre-processed. The original images, transtyped to 32-bits, are centered and reduced: their respective average intensity is subtracted and division by their standard deviation is performed. It is assumed that both signals lay close one from the other: both images are therefore combined into one to serve for synaptosomes’ detection. On each pixel, the maximum signal from both channels is retained to produce a new image, which will be subjected to both median filtering and gaussian blurring (3 pixels radius). Each potential synaptosome now appears as a bell-shaped blob, which center might be determined using a local maximum detection (tolerance to noise: 3). Second, the detections are reviewed and user-validated. Part of the original images is cropped around the local maxima and displayed to the user as a mosaic. Each thumbnail is displayed on a clickable frame, allowing the user to include or reject a synaptosome from analysis. Criteria of rejection included: presence of competing particles in the quantification area, bad focus on the particle, proximity of the image border preventing proper quantification. Finally, data is extracted, exported and displayed. A circular region is positioned over the center of the thumbnail. The centroid’s coordinates are retrieved and logged. From the two sets of coordinates (one per channel) the inter-signal distance is computed. Signal quantification is performed by placing a round region of interest (24 pixels radius) around the centroid, and measuring the integrated intensity. A measurement of the local background is performed, placing a band around the previous region. All values are logged for both channels, for all retained structures and reported in a .CSV file. Further analysis was performed using the FlowJo and GraphPad PRISM software. xy-plots of integrated intensity values are displayed with a quadrant analysis of single or double signal detections. Quadrant gates positions were defined from raw images by the experimenter. For all analyses, randomly chosen particles were displayed in a gallery to give an overview of the population analyzed (Figure 1C).

To assess the distance between two labeled dots, a plugin for ImageJ was developed. First, a binary representation of both original images was generated by a wavelet filtering algorithm (Fowler), allowing identification of the immuno-labels as individual objects. Each object was then represented by their barycenter. Two separated particles were considered associated if d < 2 µm, with d the Euclidian distance between their barycenter. To statistically determine if these associations were significant or happening by chance, we performed randomization tests. For each color channel, we fixed the position of its particle while randomizing the ones of the other channel. Since there is no underlying structure, the probability of having a particle at a certain position is identical for the whole image space. Consequently, randomization was performed by generating a complete spatial random distribution having the same number of points as the number of particles of the channel being randomized. Associations between 2 markers were then computed as explained above. The final random association values reported were defined as the mean of 10,000 randomizations.

Immunofluorescence’s data statistical analysis was performed using two-way MD ANOVA or two-way ANOVA (Table 2; Figure 2C, G, K, Figure 4B, C, D, E, F, G, Figure 5 B,D, F, H, Figure 7 B, I, Figure S4A and S5K). Distances’ data statistical analysis was performed using Kruskal-Wallis test (Figure 6A).

### Stimulated Emission Depletion (STED) Microscopy

Images were acquired using a Leica TCS SP8/STED3X microscope equipped with a HC PL APO 93X/1.30 GLYC motCORR – STED WHITE objective. We used depletion laser lines at 592nm for Alexa488 and 775nm for Alexa594 or ATTO647n fluorophores. A 25% 3D-STED effect was applied to increase Z resolution. Metrology measurements were regularly performed using fluorescent beads to test proper laser alignment. Less than 2 pixels shift between channels was measured.

### Electron Microscopy

Synaptosomes for transmission electron microscopy (Figure 1D) were fixed right after centrifugation on coverslips with a 1% Glutaraldehyde and 2% PFA (Electron Microscopy Sciences) in PBS solution and kept at 4°C until further treatment. They were then washed with PB and post-fixed in 1% osmium tetroxide and 1% K3Fe(CN)6 in PB, for 2h on ice in the dark. Washed in H_2_O and dehydrated in an ascending series of ethanol dilutions (10min in 50% ethanol, 10min in 70% ethanol, twice 15min in 95% ethanol, twice 20 min in absolute ethanol). After absolute ethanol, coverlips were lifted into Epon 812 resin (Electron Microscopy Sciences) and 50% ethanol for 2h at room temperature and then left in pure resin overnight at 4°C. Coverslips were then placed on microscope slides, embedded with capsules filled by pure resin and polymerised at 60°C for 48h. The resin block was then trimmed with razor blades. Sections, 65nm thick, were then cut using a diamond knife Ultra 35° (Diatome) with an ultra-microtome (Leica UC7) and collected on 150 mesh copper grids (Electron Microscopy Sciences).

The sections were stained with UranyLess® (Chromalys and Deltamicroscopy). Samples were then observed with an Hitachi H7650 transmission electron microscope equipped with a Gatan Orius CCD camera. Synaptosomes were identified by their size (0.5 - 2μm), their shape and the presence of intracellular compartments and organelles such as vesicles.

## Supporting information

Supp Table 4 Table S4 Proteomics comparison with Sharma et al

Supp Table 1 - Dot plots Statistics

Supp Table 2 statistical analysis

Supp Table 5 Proteomics comparison with KEGG neurotransmission pathways

Supp Table 3 Proteomic dataset

## ACKNOWLEDGMENT

Our work benefited from the excellent technical support from the central facilities at Bordeaux university: Bordeaux Imaging Center (CNRS UMS 3420, INSERM US4); Biochemistry and Biophysics of Proteins; Flow cytometry UB’FACSility (CNRS UMS 3427, INSERM US5); animal care & breeding; Genotyping; proteomics. We thank Niels Christian Danbolt for the kind gift of antibodies. We are grateful to François Georges, André Callas, Nicolas Heck and Peter Vanhoutte for fruitful discussions. Elisabeth Normand, Hajer El Oussini, Guillaume Dabee and Melissa Deshors provided an excellent support for surgeries and animal care. Julie Angibaud provided help with STED microscopy. This study benefited from the the Agence Nationale de la Recherche consortium fundings (IDEX Bordeaux ANR-10-IDEX-03-02; Labex BRAIN ANR-10-LABX-43 BRAIN; France Bio Imaging ANR-10-INBS-04) as well as funding to PT and EH (ANR-10-LABX-43 BRAIN Dolipran), funding to PT (ANR-16-CE16-0022 SynLip), and funding to TB (National Institutes of Health R01 DA018928). EH received funding from Fondation pour la Recherche Médicale (ING20150532192) and EH and DP from ANR (DopamineHub-19-CE16-0003-01) and from Conseil Régional de Nouvelle Aquitaine (ParkSynGref). RW holds a fellowship from the French ministry of research.

## AUTHOR CONTRIBUTIONS

Experimental design: DP, EH, MEP, MPr, PT, VPB; Experimental work: EH, MEP, MPr, MFA, PL, VPB; Technical contribution to experiments: CM, FPC, FL, MFA, MPe, RW, SC, SL, VDP, VP; Software programing and data analysis: EH, FL, FPC, MEP, MPr, PL, SC; Reagent sharing: TB; Funding : DP, EH, PT; Article writing: DP, EH, MEP, MP, PL, PT, VPB; Article editing: all authors

## CONFLICT OF INTEREST

The authors declare no conflict of interest

---

**Figure S1:**
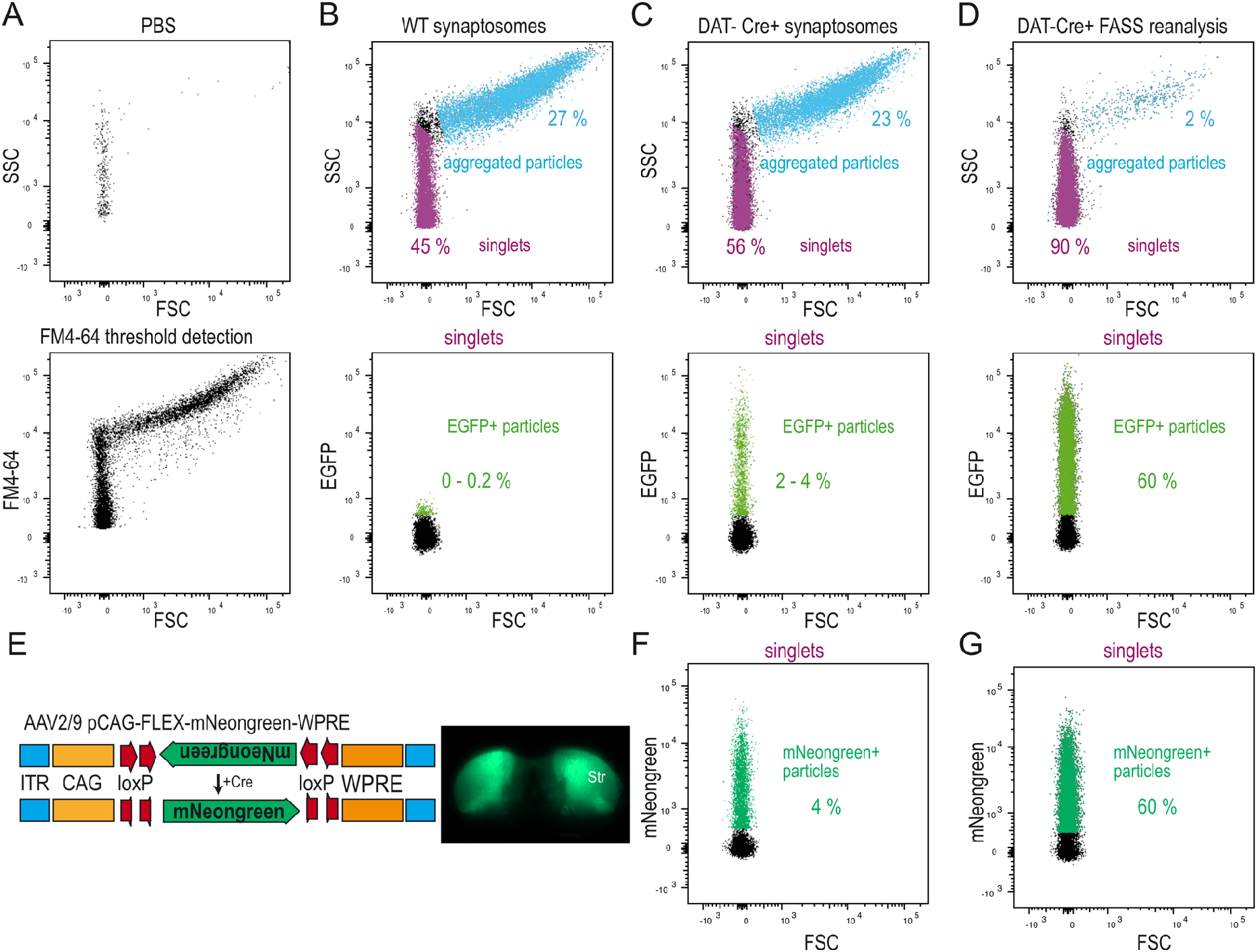
DA-FASS gating strategy. **(A-D)** Representative FASS gate settings and particles detection for DAT-Cre EGFP+ synaptosomes sorting. **(A)** Analysis of PBS allows to define background noise of the thresholding using FM4-64 lipopholic styryl dye used (top). The noise is less than 500 events per minute. Particle detection using FM4-64 thresholding (bottom). **(B)** Side scatter (SSC) and forward scatter (FSC) analysis of synaptosomes allows defining aggregated particles (27%, light blue gate) and singlets (45%, magenta gate; top). Singlets gate was defined experimentally through trials and error as published previously (Luquet et al., 2016). Singlets are further analysed for EGFP fluorescence (bottom). Saline synaptosomes display low autofluorescence. **(C)** DAT-Cre+ synaptosomes samples showed 23% of aggregated particles and 56% of singlets on this example (top). 2-4% of the singlets were significantly fluorescent in the EGFP channel. **(D)** Particles gated as “singlets” and “EGFP+” were sorted and re-analysed showing a drop in the proportion of aggregated particles (2%) and a steep rise in singlets (90%; top). Up to 60% of singlets were EGFP+ (bottom). **(E)** (Left) Cre-dependent AAV expressing mNeongreen. (Right) mNeongreen striatal expression. **(F)** Before sorting 4% of the singlets were significantly fluorescent in the mNeongreen channel. **(G)** Same as D bottom, after sorting up to 60% of singlets were mNeongreen+ in the DA-FASS sample.

**Figure S2:**
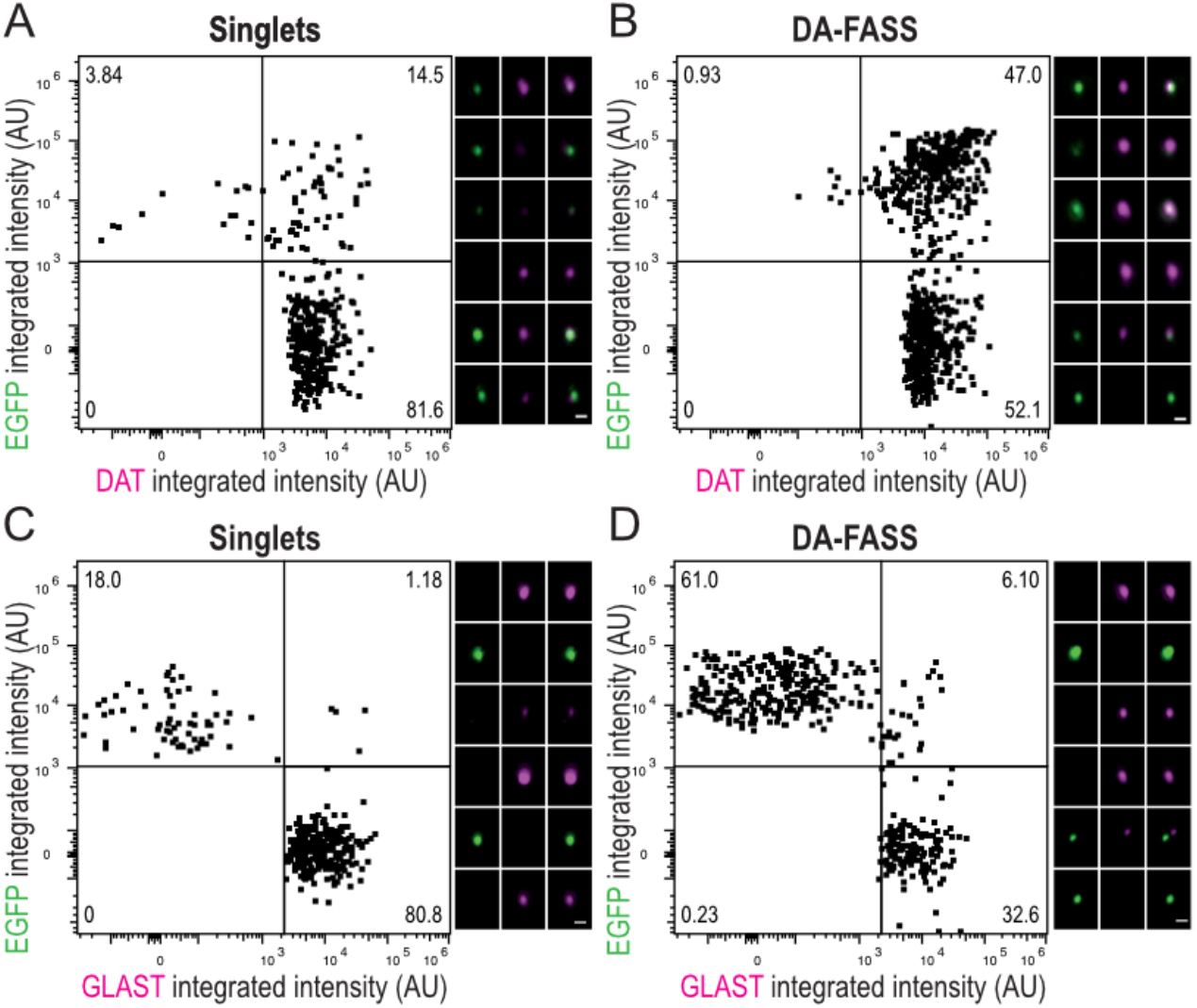
Immunofluorescence of DA-FASS synaptosomes. **(A-B)** (Left) Dot plots of singlets and DA-FASS synaptosomes stained for the dopamine transporter DAT (x-axis) and EGFP (y-axis). (Right) Galleries of representative epifluorescence images of individual synaptosomes. Population of particles positive for both EGFP and DAT (upper right quadrant) increases from 15% in the singlets to 47% in the DA-FASS sample. **(C-D)** Dot plots of intensity signal of singlets and DA-FASS synaptosomes stained for EGFP and the astrocyte membrane marker Slc1a3/GLAST and galleries of representative images. Note the very low representation of double positive particles. Scale bar = 1 μm.

**Figure S3:**
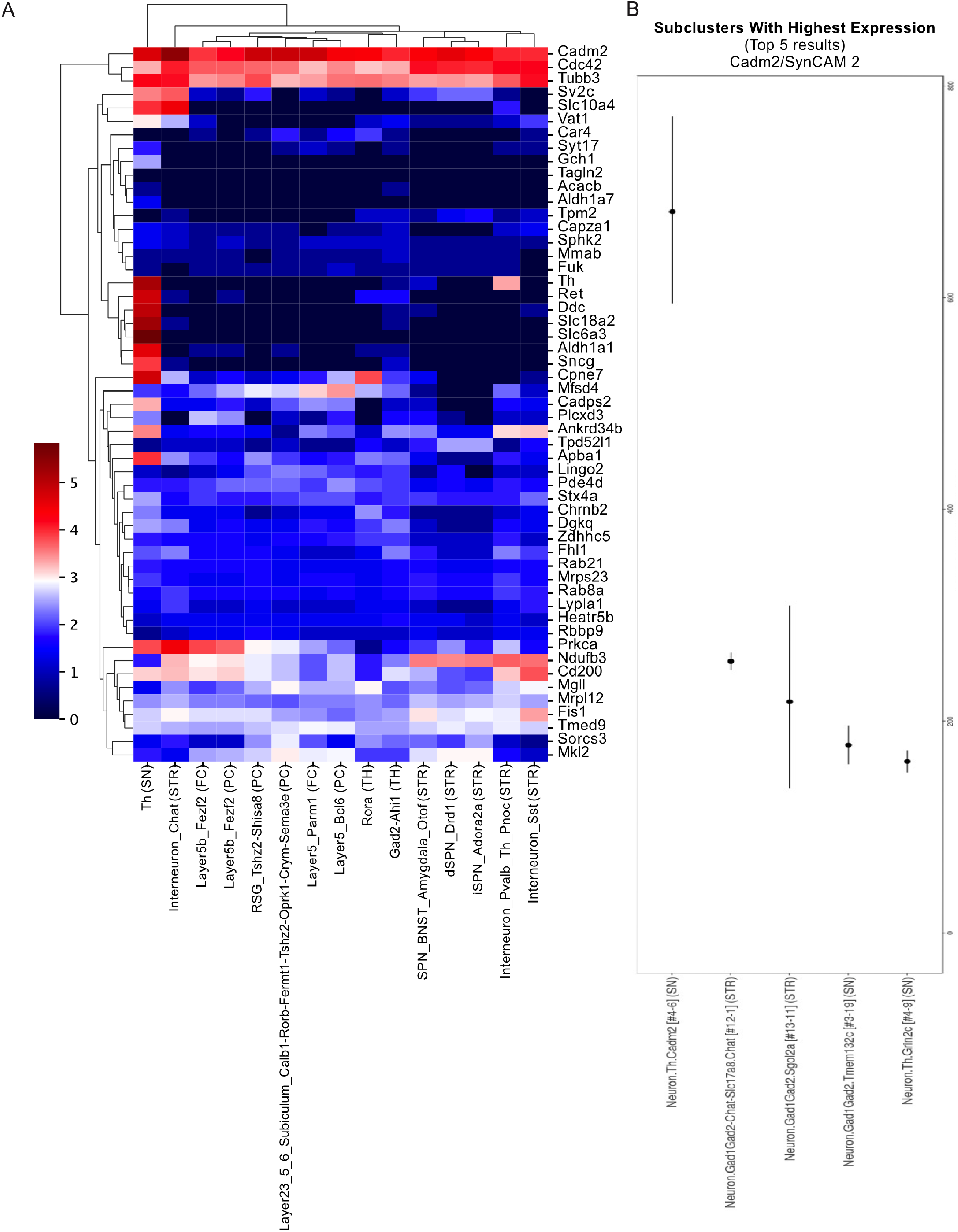
Meta-analysis of enriched proteins with single cell RNA databases. **(A)** Extended Heatmap showing cell type specific mRNA abundance in neuronal cells from or projecting to the striatum (STR; Substancia Nigra, SN; Thalamus, TH; Frontal Cortex, FC; Posterior Cortex, PC) of 53 of the enriched DA-FASS proteins present in the DropViz single cell RNA sequencing database (Stx4a = Stx4, Mfsd4 = Mfsd4a, Fuk = Fcsk, Car4 = Ca4, Mkl2 = Mrtfb) (DropViz; (Saunders *et al*, 2018)). **(B)** Drop Viz representation of Cadm2/ SynCAM 2 brain cellular subclusters with highest expression. Data represented per 100 000 transcripts in sub cluster. Note that the highest Cadm2/SynCAM 2 mRNA expression throughout the brain is found in a sub cluster of substantia nigra Th neurons.

**Figure S4:**
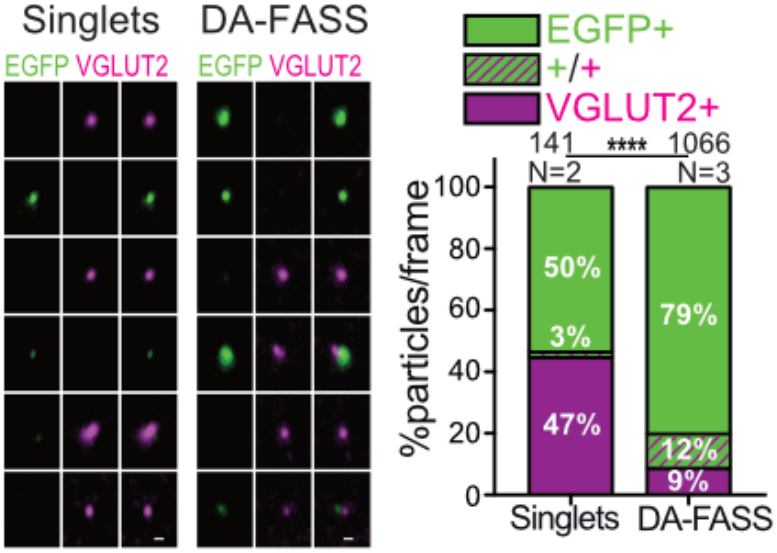
Characterization of thalamo-striatal hub synapses. Epifluorescence images (left) of a representative sample of synaptosome populations (singlets and DA FASS) labelled with anti-EGFP and anti-VGLUT2. labelling mainly thalamo-striatal terminals. (Right) Analysis of particle proportions per frame. n=6, for singlets and n=14 for FASS samples. The EGFP+/VGLUT2+ population increases from 3% to 12%. Data represented as mean, ****p<0.001. Two-way MD ANOVA.

**Figure S5:**
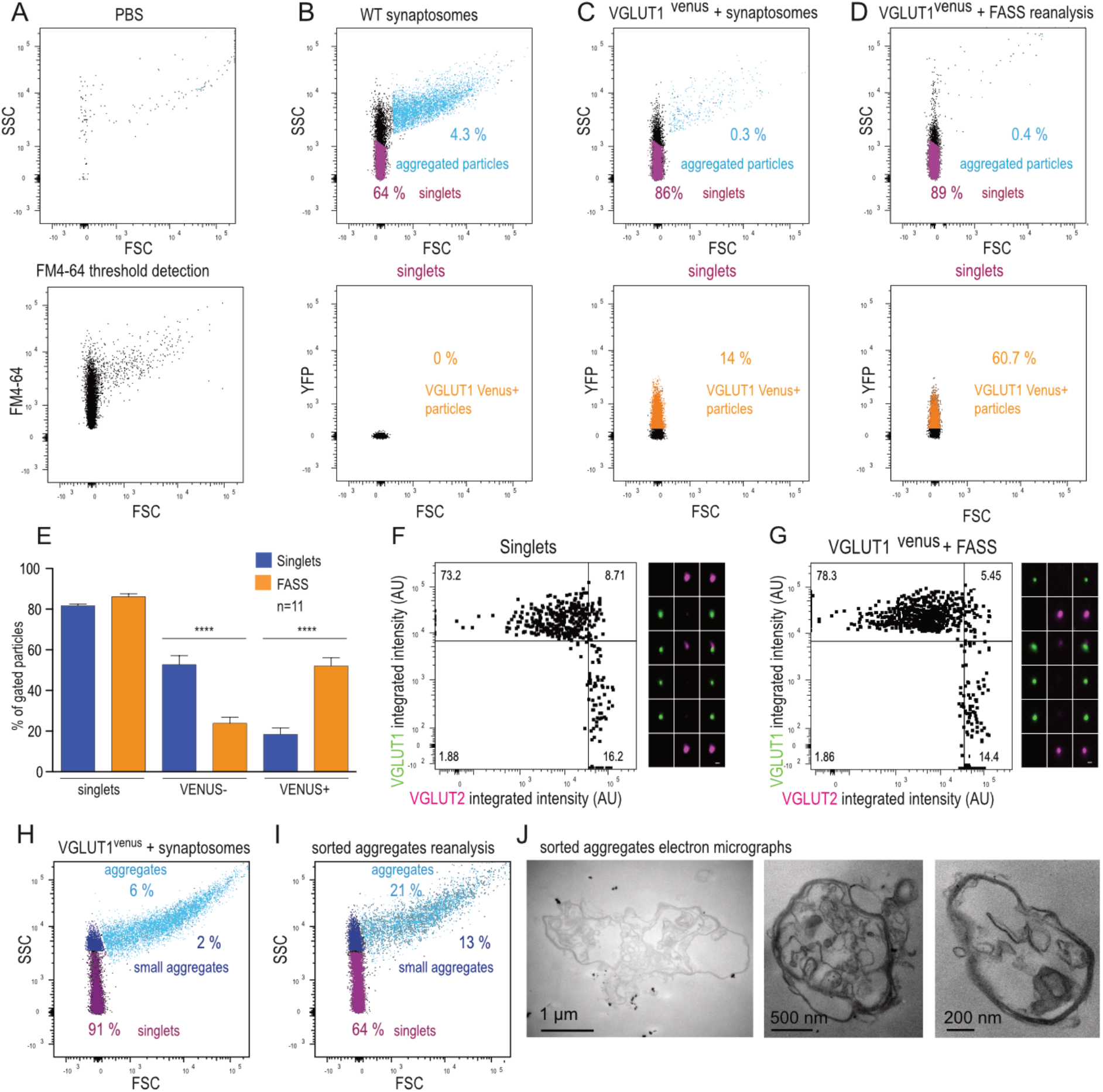
Positive and negative controls to dopamine hub synapse identification. **(A-D)** VGLUT1^venus^ FASS was performed to test for the non specific aggregation of VGLUT1 cortico-striatal synapses with VGLUT1 thalamo-striatal synapses. Also we tested for the association of VGLUT1^venus^ synapses with Th varicosities in this alternative FASS protocol (see Figure 7). **(A)** Analysis of a PBS sample was used to define background noise in FM4-64 lipophilic styryl dye used for thresholding (top). The noise was less than 500 events per minute. Synaptosome sample detection using FM4-64 thresholding (bottom). **(B)** WT synaptosomes display aggregated particles (4.3%, light blue) and singlets (64%, magenta). Singlets gate was defined experimentally through trials and error as published previously (Luquet et al., 2017). Singlets were further analysed for Venus fluorescence to determine the autofluorescence level. **(C)** Singlets sorted VGLUT1^venus^ synaptosomes samples showed 0.3% of aggregated particles and 86% of singlets. 14% of synaptosomes were detected in the venus gate. **(D)** Sorted VGLUT1^venus^ “singlets” were re-analysed. VGLUT1^venus^ + synaptosomes displayed 0.4% of aggregated particles and 89% singlets. Up to 60.7% of sorted singlets were indeed detected in the venus gate. **(E)** Average striatal VGLUT1^venus^ FASS results. Singlets (blue) and FASS (orange) samples for the different gates: singlets, VENUS-, VENUS+. Note the steep increase in VENUS+ particles and significant decrease in VENUS-contaminants through the FASS process. n=11 different sorts/condition, all data are mean ±SEM. Post-Hoc ****p < 0.001. two-way MD ANOVA. **(F-G)** Dot plots singlets and VGLUT1^venus^ + FASS synaptosomes stained for VGLUT1 and VGLUT2. and galleries of representative epifluorescence images. Note the low representation of synaptosomes associating VGLUT1 (cortico-striatal inputs) and VGLUT2 (Thalamo-striatal inputs). Indeed VGLUT1/-2 double positives do not enrich through FASS. **(H-I)** Representative FASS gating for aggregates sorting. (H) Sucrose/Ficoll VGLUT1^venus^ synaptosome samples showed 6% of aggregated particles (light blue). (I) Particles gated as “aggregates” and large tissue fragments were sorted and reanalysed. Small and large aggregates represented 34% of particles after sorting. Indeed unspecific aggregates tend to break down when facing shearing forces exisiting at the nozzle of the FACS and generates singlets at reanalysis. **(J)** Electron micrographs of sorted aggregated particles. Aggregates appear much larger than hub synapses. Their cellular content is difficult to identify though myelin membranes may be recognized on some of them. Scale bar, 1μm, 500nm, 200nm, (from left to right).

**Figure S6:**
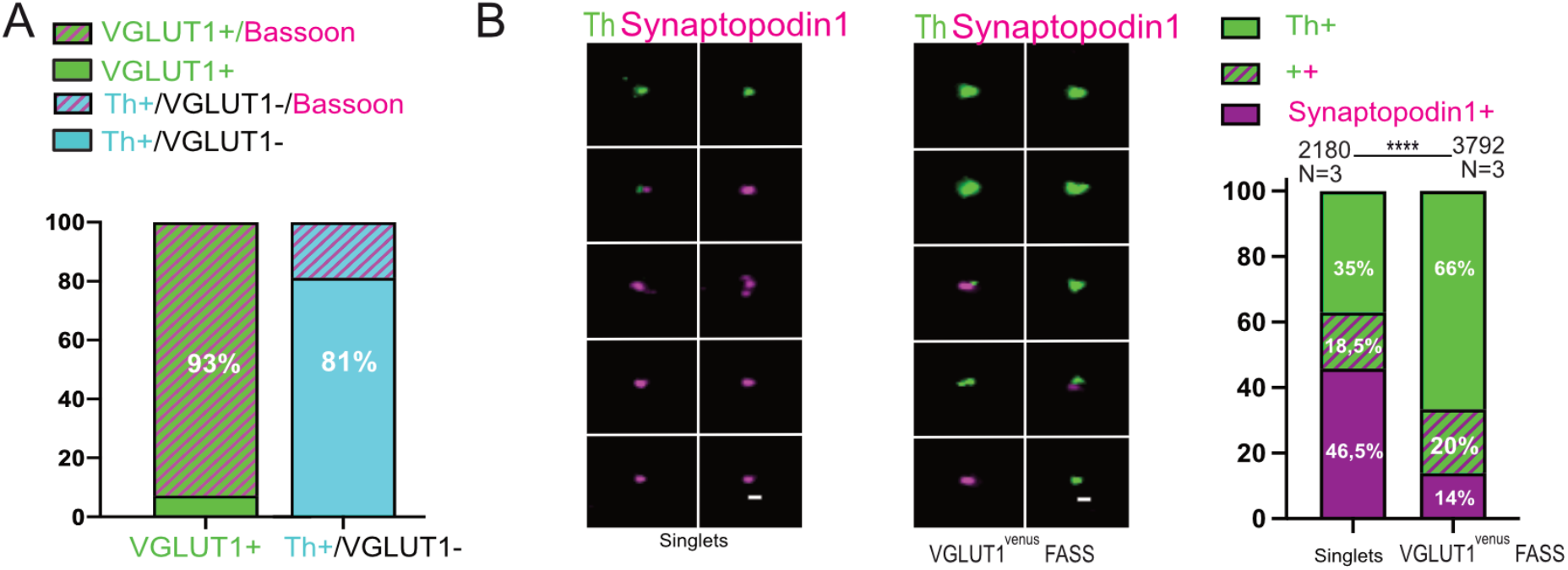
Characterization of VGLUT1/Th hub synapses. **(A)** Proportion of Bassoon in VGLUT1+ or Th+/VGLUT1-synaptosomes. **(B)** Epifluorescence images (left) of a representative sample of synaptosome populations (singlets and VGLUT1^Venus^ + FASS) labelled with anti-Th (green) and anti-Synaptopodin1 (magenta). (Right) Analysis of Th and Synaptopodin 1 particle proportions per frame. n=10, for singlets and n=11 for FASS samples. Data represented as mean, interaction ****p<0.001. Two-way MD ANOVA.

## Notes

### Competing Interest Statement

The authors have declared no competing interest.

### Summary of Updates

Major change with new datasets and new authors.

