## Supplementary material for "Synaptomic analysis of dopaminergic inputs reveal hub synapses in the mouse striatum": Supp Table 1 - Dot plots Statistics

| **Fig. 2 A,B** | **Quadrant** | **Singlets (n=3)** | **FASS (n=3)** |
| --- | --- | --- | --- |
|  | n. of total particles | 298 | 2378 |
|  | EGFP+/TH - | 11.1 % | 6.18 % |
|  | EGFP-/ TH + | 33.9 % | 12.1 % |
|  | EGFP+/TH + | 54.0 % | 81.7 % |
| **Fig. 2 E,F** | **Quadrant** | **Unsorted (n=2)** | **FASS (n=2)** |
|  | n. of total particles | 1152 | 1944 |
|  | EGFP+/D1R - | 14.3 % | 26.1 % |
|  | EGFP-/ D1R + | 81.0 % | 42.0 % |
|  | EGFP+/D1R + | 3.99 % | 31.8 % |
| **Fig. 2 I,J** | **Quadrant** | **Unsorted (n=3)** | **FASS (n=3)** |
|  | n. of total particles |  |  |
|  | EGFP+/D2R - | 6.47 % | 12.3 % |
|  | EGFP-/ D2R + | 76.0 % | 37.5 % |
|  | EGFP+/D2R + | 13.7 % | 50.0 % |
| **Fig.S2 A,B** | **Quadrant** | **Unsorted (n=5)** | **FASS (n=5)** |
|  | n. of total particles | 365 | 751 |
|  | EGFP+/DAT - | 3.84 % | 0.93 % |
|  | EGFP-/ DAT + | 81.6 % | 47.0 % |
|  | EGFP+/DAT + | 14.5 % | 52.1 % |
| **Fig.S2 C,D** | **Quadrant** | **Unsorted (n=1)** | **FASS (n=1)** |
|  | n. of total particles | 338 | 426 |
|  | EGFP+/GLAST - | 18.0 % | 61.0 % |
|  | EGFP-/ GLAST + | 80.8 % | 32.6 % |
|  | EGFP+/GLAST + | 1.18 % | 6.10 % |
