## Supplementary material for "Synaptomic analysis of dopaminergic inputs reveal hub synapses in the mouse striatum": Supp Table 2 statistical analysis

| Figure | Labelling | Condition | Mean ± SEM | Source of Variation | F (DFn, DFd) | p value | Post-Hoc (Šídák) | p value | Test |
| --- | --- | --- | --- | --- | --- | --- | --- | --- | --- |
| Fig. 1D | TH | H | 1 ± 0,26 n=3 | Fraction x Protein | F (8, 18) = 0,5759 | p=0,7844 |  |  | Two-way  RM ANOVA |
|  |  | P1 | 0,36 ± 0,07 n=3 | Fraction | F (8, 18) = 4,630 | **p=0,0033 | |  |  |
|  |  | S2 | 0,77 ± 0,1 n=3 | Protein | F (1, 18) = 5,030 | *p=0,0377 |  |  |  |
|  |  | P2 | 1,2 ± 0,27 n=3 |  |  |  |  |  |  |
|  |  | Syn | 1,5 ± 0,31 n=3 |  |  |  |  |  |  |
|  |  | LS1 | 2,2 ± 0,19 n=3 |  |  |  |  |  |  |
|  |  | SPM | 0,26 ± 0,02 n=3 |  |  |  |  |  |  |
|  |  | LS2 | 1,58 ± 0,32 n=3 |  |  |  |  |  |  |
|  |  | LP2 | 0,82 ± 0,4 n=3 |  |  |  |  |  |  |
|  | GFP | H | 1 ± 0,53 n=3 |  |  |  |  |  |  |
|  |  | P1 | 0,23 ± 0,09 n=3 |  |  |  |  |  |  |
|  |  | S2 | 0,49 ± 0,23 n=3 |  |  |  |  |  |  |
|  |  | P2 | 1,12 ± 0,52 n=3 |  |  |  |  |  |  |
|  |  | Syn | 1,48 ± 0,71 n=3 |  |  |  |  |  |  |
|  |  | LS1 | 1,8 ± 0,5 n=3 |  |  |  |  |  |  |
|  |  | SPM | 0,07 ± 0,02 n=3 |  |  |  |  |  |  |
|  |  | LS2 | 1,29 ± 0,22 n=3 |  |  |  |  |  |  |
|  |  | LP2 | 0,03 ± 0,01 n=3 |  |  |  |  |  |  |
| Fig. 1F | Singlets | Singlets | 73,79 ± 4,61 n=9 | Interaction | F (2, 45) = 88,43 | ****p<0,0001 | Singlets - FASS |  | Two-way  MD ANOVA |
|  |  | FASS | 83,91 ± 1,25 n=8 | Condition | F (1, 45) = 7,275 | **p=0,0098 | Singlets | p=0,0657 |  |
|  |  |  |  | Fluorescence | F (2, 45) = 151,4 | ****p<0,0001 | EGPP- | ****p<0,0001 | |
|  | EGPP- | Singlets | 66,14 ± 3,97 n=9 | |  |  | EGFP+ | ****p<0,0001 | |
|  |  | FASS | 30,94 ± 2,75 n=8 | |  |  |  |  |  |
|  | EGFP+ | Singlets | 3,86 ± 0,52 n=9 |  |  |  |  |  |  |
|  |  | FASS | 48,94 ± 2,3 n=8 |  |  |  |  |  |  |
| Fig. 2C | TH+ | Singlets | 25,75 ± 4,18 n=14 | Interaction | F (2, 99) = 49,29 | ****p<0,0001 | Singlets - FASS |  | Two-way  MD ANOVA |
|  |  | FASS | 8,18 ± 0,76 n=21 | Condition | F (1, 99) = 0,03 | p=0,8562 | TH+ | ****p<0,0001 | |
|  |  |  |  | Staining | F (2, 99) = 414,5 | ****p<0,0001 | ++ | ****p<0,0001 | |
|  | ++ | Singlets | 56,84 ± 2,26 n=14 | |  |  | EGFP + | p=0,2179 |  |
|  |  | FASS | 80,76 ± 1,53 n=21 | |  |  |  |  |  |
|  | EGFP+ | Singlets | 16,39 ± 2,5 n=14 | |  |  |  |  |  |
|  |  | FASS | 10,99 ± 1,69 n=21 | |  |  |  |  |  |
| Figure | Labelling | Condition | Mean ± SEM | Source of Variation | F (DFn, DFd) | p value | Post-Hoc (Šídák) | p value | Test |
| Fig. 2G | D1R+ | Singlets | 86,61 ± 1,01 n=20 | Interaction | F (2, 120) = 256,0 | ****p<0,0001 | Singlets - FASS |  | Two-way  MD ANOVA |
|  |  | FASS | 41,68 ± 1,79 n=22 | Condition | F (1, 120) = 0,01 | p=0,9182 | D1R+ | ****p<0,0001 | |
|  |  |  |  | Staining | F (2, 120) = 474,0 | ****p<0,0001 | ++ | ****p<0,0001 | |
|  | ++ | Singlets | 4,6 ± 0,77 n=20 |  |  |  | EGFP+ | ****p<0,0001 | |
|  |  | FASS | 33,61 ± 2,1 n=22 | |  |  |  |  |  |
|  | EGFP+ | Singlets | 8,79 ± 1,02 n=20 | |  |  |  |  |  |
|  |  | FASS | 24,26 ± 2,59 n=22 | |  |  |  |  |  |
| Fig. 2K | D2R+ | Singlets | 32,15 ± 5,28 n=32 | Interaction | F (2, 180) = 20,78 | ****p<0,0001 | Singlets - FASS |  | Two-way  MD ANOVA |
|  |  | FASS | 8,99 ± 0,93 n=30 | Condition | F (1, 180) = 0,01 | p=0,9043 | D2R+ | ****p<0,0001 | |
|  |  |  |  | Staining | F (2, 180) = 131,8 | ****p<0,0001 | ++ | ****p<0,0001 | |
|  | ++ | Singlets | 55,05 ± 6,01 n=32 | |  |  | EGFP+ | p=0,9939 |  |
|  |  | FASS | 78,09 ± 1,34 n=30 | |  |  |  |  |  |
|  | EGFP+ | Singlets | 12,08 ± 2,03 n=32 | |  |  |  |  |  |
|  |  | FASS | 13,25 ± 1,4 n=30 | |  |  |  |  |  |
| Fig. 4B | CPNE7+ | Singlets | 42,61 ± 2,4 n=31 | Interaction | F (2, 180) = 131,9 | ****p<0,0001 | Singlets - FASS |  | Two-way  MD ANOVA |
|  |  | FASS | 16,21 ± 1,28 n=31 | Condition | F (1, 180) = 0,0004 | p=0,9841 | CPNE7+ | ****p<0,0001 | |
|  |  |  |  | Staining | F (2, 180) = 570,4 | ****p<0,0001 | ++ | p=0,9951 |  |
|  | ++ | Singlets | 8 ± 0,88 n=31 |  |  |  | TH+ | ****p<0,0001 | |
|  |  | FASS | 8,49 ± 0,44 n=31 | |  |  |  |  |  |
|  | TH+ | Singlets | 49,3 ± 2,35 n=31 | |  |  |  |  |  |
|  |  | FASS | 75,29 ± 1,32 n=31 | |  |  |  |  |  |
| Fig. 4C | Mint1+ | Singlets | 66,14 ± 3,09 n=30 | Interaction | F (2, 183) = 163,7 | ****p<0,0001 | Singlets - FASS |  | Two-way  MD ANOVA |
|  |  | FASS | 29,92 ± 1,27 n=33 | Condition | F (1, 183) = 0,0009 | p=0,9754 | Mint1+ | ****p<0,0001 | |
|  |  |  |  | Staining | F (2, 183) = 316,5 | ****p<0,0001 | ++ | p=0,2048 |  |
|  | ++ | Singlets | 3,56 ± 0,5 n=30 |  |  |  | TH+ | ****p<0,0001 | |
|  |  | FASS | 8,35 ± 0,65 n=33 | |  |  |  |  |  |
|  | TH+ | Singlets | 30,05 ± 2,99 n=30 | |  |  |  |  |  |
|  |  | FASS | 61,34 ± 1,33 n=33 | |  |  |  |  |  |
| Fig. 4D | CADPS2+ | Singlets | 57,71 ± 3,33 n=30 | Interaction | F (2, 187) = 110,5 | ****p<0,0001 | Singlets - FASS |  | Two-way  MD ANOVA |
|  |  | FASS | 22,63 ± 1,1 n=34 | Condition | F (1, 187) = 0,004 | p=0,9511 | CADPS2+ | ****p<0,0001 | |
|  |  |  |  | Staining | F (2, 187) = 88,12 | ****p<0,0001 | +/+ | *p=0,0434 |  |
|  | ++ | Singlets | 12,99 ± 1,65 n=30 | |  |  | TH+ | ****p<0,0001 | |
|  |  | FASS | 20,54 ± 1,49 n=34 | |  |  |  |  |  |
|  | TH+ | Singlets | 28,97 ± 3,41 n=31 | |  |  |  |  |  |
|  |  | FASS | 56,83 ± 1,11 n=34 | |  |  |  |  |  |
| Figure | Labelling | Condition | Mean ± SEM | Source of Variation | F (DFn, DFd) | p value | Post-Hoc (Šídák) | p value | Test |
| Fig. 4E | SynCAM2+ | Singlets | 53,18 ± 2,38 n=12 | Interaction | F (2, 60) = 84,92 | ****p<0,0001 | Singlets - FASS |  | Two-way  MD ANOVA |
|  |  | FASS | 18,87 ± 3,05 n=10 | Condition | F (1, 60) = 4,371e-005 | p=0,9947 | SynCAM2+ | ****p<0,0001 | |
|  |  |  |  | Staining | F (2, 60) = 73,75 | ****p<0,0001 | ++ | ****p<0,0001 | |
|  | ++ | Singlets | 28,69 ± 1,56 n=12 | |  |  | EGFP+ | p=0,1265 |  |
|  |  | FASS | 71,76 ± 5,67 n=10 | |  |  |  |  |  |
|  | EGFP+ | Singlets | 18,18 ± 1,93 n=12 | |  |  |  |  |  |
|  |  | FASS | 9,37 ± 2,81 n=10 | |  |  |  |  |  |
| Fig. 4F | Stx4+ | Singlets | 62,45 ± 3,11 n=31 | Interaction | F (2, 177) = 69,25 | ****p<0,0001 | Singlets - FASS |  | Two-way  MD ANOVA |
|  |  | FASS | 29,18 ± 1,74 n=30 | Condition | F (1, 177) = 0,003998 | p=0,9497 | Stx4+ | ****p<0,0001 | |
|  |  |  |  | Staining | F (2, 177) = 49,15 | ****p<0,0001 | ++ | ****p<0,0001 | |
|  | ++ | Singlets | 7,16 ± 1,22 n=31 | |  |  | TH+ | p=0,1185 |  |
|  |  | FASS | 33,31 ± 3,08 n=30 | |  |  |  |  |  |
|  | TH+ | Singlets | 30 ± 2,66 n=31 |  |  |  |  |  |  |
|  |  | FASS | 37,51 ± 3,06 n=30 | |  |  |  |  |  |
| Fig. 4G | Mgll+ | Singlets | 31,18 ± 2,17 n=33 | Interaction | F (2, 192) = 98,57 | ****p<0,0001 | Singlets - FASS |  | Two-way  MD ANOVA |
|  |  | FASS | 8,83 ± 0,57 n=33 | Condition | F (1, 192) = 4,217e-005 | p=0,9948 | Mgll+ | ****p<0,0001 | |
|  |  |  |  | Staining | F (2, 192) = 1242 | ****p<0,0001 | ++ | **p=0,0015 |  |
|  | ++ | Singlets | 3,22 ± 0,54 n=33 | |  |  | TH+ | ****p<0,0001 | |
|  |  | FASS | 10,28 ± 0,95 n=33 | |  |  |  |  |  |
|  | TH+ | Singlets | 65,48 ± 2,08 n=33 | |  |  |  |  |  |
|  |  | FASS | 80,79 ± 1,17 n=33 | |  |  |  |  |  |
| Figure | Labelling | Condition | Mean ± SEM | Source of Variation | F (DFn, DFd) | p value | Post-Hoc (Šídák) | p value | Test |
| Fig. 5B | Synapsin+ | Singlets | 83,2 ± 1,06 n=9 | Interaction | F (2, 51) = 237,8 | ****p<0,0001 | Singlets - FASS |  | Two-way  MD ANOVA |
|  |  | FASS | 33,78 ± 2,38 n=10 | Condition | F (1, 51) = 0,01020 | p=0,9199 | Synapsin+ | ****p<0,0001 | |
|  |  |  |  | Staining | F (2, 51) = 237,5 | ****p<0,0001 | ++ | ****p<0,0001 | |
|  | ++ | Singlets | 6,62 ± 1,15 n=9 |  |  |  | EGFP+ | ***p=0,0006 | |
|  |  | FASS | 44,88 ± 3,31 n=10 | |  |  |  |  |  |
|  | EGFP+ | Singlets | 9,64 ± 1,43 n=9 |  |  |  |  |  |  |
|  |  | FASS | 21,31 ± 1,6 n=10 | |  |  |  |  |  |
| Fig. 5D | VGLUT1+ | Singlets | 76,23 ± 1,62 n=12 | Interaction | F (2, 63) = 91,49 | ****p<0,0001 | Singlets - FASS |  | Two-way  MD ANOVA |
|  |  | FASS | 30,93 ± 4,53 n=11 | Condition | F (1, 63) = 0,001 | p=0,9695 | Vglut1 only | ****p<0,0001 | |
|  |  |  |  | Staining | F (2, 63) = 92,06 | ****p<0,0001 | ++ | **p=0,0022 |  |
|  | ++ | Singlets | 5,92 ± 1,39 n=12 | |  |  | EGFP only | ****p<0,0001 | |
|  |  | FASS | 20,78 ± 2,68 n=11 | |  |  |  |  |  |
|  | EGFP+ | Singlets | 17,8 ± 1,68 n=12 | |  |  |  |  |  |
|  |  | FASS | 48,52 ± 4,61 n=11 | |  |  |  |  |  |
| Fig. 5F | VIAAT+ | Singlets | 68,19 ± 3,24 n=9 | Interaction | F (2, 78) = 54,90 | ****p<0,0001 | Singlets - FASS |  | Two-way  MD ANOVA |
|  |  | FASS | 25,93 ± 1,9 n=19 | Condition | F (1, 78) = 0,04 | p=0,8438 | VIAAT+ | ****p<0,0001 | |
|  |  |  |  | Staining | F (2, 78) = 55,34 | ****p<0,0001 | ++ | *p=0,0158 |  |
|  | ++ | Singlets | 4,11 ± 0,92 n=9 |  |  |  | EGFP+ | ****p<0,0001 | |
|  |  | FASS | 18,76 ± 3,22 n=19 | |  |  |  |  |  |
|  | EGFP+ | Singlets | 25,96 ± 2,72 n=9 | |  |  |  |  |  |
|  |  | FASS | 55,32 ± 4,28 n=19 | |  |  |  |  |  |
| Fig. 5H | VAChT+ | Singlets | 62,5 ± 2,08 n=30 | Interaction | F (2, 144) = 180,3 | ****p<0,0001 | Singlets - FASS |  | Two-way  MD ANOVA |
|  |  | FASS | 28,01 ± 1,34 n=20 | Condition | F (1, 144) = 0,01571 | p=0,9004 | VAChT+ | ****p<0,0001 | |
|  |  |  |  | Staining | F (2, 144) = 412,2 | ****p<0,0001 | ++ | **p=0,0011 |  |
|  | ++ | Singlets | 1,68 ± 0,37 n=30 | |  |  | TH+ | ****p<0,0001 | |
|  |  | FASS | 10,19 ± 0,65 n=20 | |  |  |  |  |  |
|  |  |  | ± n= |  |  |  |  |  |  |
|  | TH+ | Singlets | 35,23 ± 2,18 n=30 | |  |  |  |  |  |
|  |  | FASS | 61,71 ± 1,19 n=20 | |  |  |  |  |  |
| Figure | Labelling | Condition | Mean ± SEM | Source of Variation | F (DFn, DFd) | p value | Post-Hoc (Šídák) | p value | Test |
| Fig. 6A | TH | FASS | 0,25 ± 0,01 n=971 | |  | ****p<0,0001 | TH-Staining |  | Kruskal-  Wallis |
|  | SynCAM2 | FASS | 0,32 ± 0,01 n=1010 | |  |  | TH vs. SynCAM2 | ****p<0,0001 | |
|  | D2R | FASS | 0,33 ± 0,01 n=471 | |  |  | TH vs. D2R | ****p<0,0001 | |
|  | D1R | FASS | 0,35 ± 0,01 n=184 | |  |  | TH vs. D1R | ****p<0,0001 | |
|  | Synapsin | FASS | 0,43 ± 0,01 n=521 | |  |  | TH vs. Synapsin | ****p<0,0001 | |
|  | VAChT | FASS | 0,51 ± 0,02 n=217 | |  |  | TH vs. VAChT | ****p<0,0001 | |
|  | VIAAT | FASS | 0,56 ± 0,02 n=337 | |  |  | TH vs. VIAAT | ****p<0,0001 | |
|  | VGLUT2 | FASS | 0,62 ± 0,03 n=117 | |  |  | TH vs. VGLUT2 | ****p<0,0001 | |
|  | VGLUT1 | FASS | 0,64 ± 0,02 n=231 | |  |  | TH vs. VGLUT1 | ****p<0,0001 | |
| Fig. 7B | TH+ | Singlets | 55,06 ± 3,19 n=19 | Interaction | F (2, 111) = 47,14 | ****p<0,0001 | Singlets - FASS |  | Two-way  MD ANOVA |
|  |  | FASS | 22,03 ± 1,55 n=20 | Condition | F (1, 111) = 0,002573 | p=0,9596 | TH+ | ****p<0,0001 | |
|  |  |  |  | Staining | F (2, 111) = 15,54 | ****p<0,0001 | ++ | ***p=0,0006 | |
|  | ++ | Singlets | 15,46 ± 2,6 n=19 | |  |  | VGLUT1+ | ***p=0,0002 | |
|  |  | FASS | 31,5 ± 4,02 n=20 | |  |  |  |  |  |
|  | VGLUT1+ | Singlets | 28,14 ± 2,6 n=19 | |  |  |  |  |  |
|  |  | FASS | 45,5 ± 3,16 n=20 | |  |  |  |  |  |
| Fig. 7C | TH | Singlets | 437200 ± 79456,71 n=3 | Fraction x Protein | F (1, 4) = 3,928 | p=0,1185 |  |  | Two-way  RM ANOVA |
|  |  | FASS | 357648 ± 107551,79 n=3 | Fraction | F (1, 4) = 1,861 | p=0,2442 |  |  |  |
|  |  |  |  | Protein | F (1, 4) = 2,193 | p=0,2128 |  |  |  |
|  | VENUS | Singlets | 391944,33 ± 60156,53 n=3 | | |  |  |  |  |
|  |  | FASS | 670425,33 ± 73824,66 n=3 | | |  |  |  |  |
| Figure | Labelling | Condition | Mean ± SEM | Source of Variation | F (DFn, DFd) | p value | Post-Hoc (Šídák) | p value | Test |
| Fig. 7D | VGLUT1+ |  |  |  |  |  |  |  |  |
|  |  | TH- | 143489 ± 2037 n=3609 | |  | ****p<0,0001 | |  | Mann Whitney |
|  |  | TH+ | 182796 ± 4513 n=1206 | |  | ****p<0,0001 | |  | Kolmogorov-  Smirnov |
|  | Bassoon+ |  |  |  |  |  |  |  |  |
|  |  | TH- | 138149 ± 1856 n=3609 | |  | ****p<0,0001 | |  | Mann Whitney |
|  |  | TH+ | 205934 ± 7061 n=1206 | |  | ****p<0,0001 | |  | Kolmogorov-  Smirnov |
| Fig. 7F |  |  |  |  |  |  |  |  |  |
|  | Homer+ |  |  |  |  |  |  |  |  |
|  |  | TH- | 55356 ± 1052 n=1877 | |  | ****p<0,0001 | |  | Mann Whitney |
|  |  | TH+ | 64756 ± 1385 n=1536 | |  | ****p<0,0001 | |  | Kolmogorov-  Smirnov |
| Fig. 7G |  |  |  |  |  |  |  |  |  |
|  | PSD95+ |  |  |  |  |  |  |  |  |
|  |  | TH- | 21758 ± 471,1 n=2232 | |  | ****p<0,0001 | |  | Mann Whitney |
|  |  | TH+ | 20149 ± 580,9 n=1533 | |  | ***p=0,0004 | |  | Kolmogorov-  Smirnov |
| Fig. 7H | Synaptopodin1+ |  |  |  |  |  |  |  |  |
|  |  | TH- | 213786 ± 6158 n=1528 | |  | **p=0,0015 | |  | Mann Whitney |
|  |  | TH+ | 382663 ± 36567 n=725 | |  | *p=0,0272 |  |  | Kolmogorov-  Smirnov |
| Fig. 7I | VGLUT1+ | TH- | 68,71 ± 1,17 n=81 | Interaction | F (1, 320) = 95,48 | ****p<0,0001 | TH- - TH+ |  | Two-way  ANOVA |
|  |  | TH+ | 54,63 ± 1,7 n=81 | Row Factor | F (1, 320) = 0,1688 | p=0,6815 | VGLUT1+ | ****p<0,0001 | |
|  |  |  |  | Column Factor | F (1, 320) = 326,0 | ****p<0,0001 | VGLUT1+ Synaptopodin1+ | ****p<0,0001 | |
|  | VGLUT1+ Synaptopodin1+ | TH- | 30,22 ± 0,92 n=81 | |  |  |  |  |  |
|  |  | TH+ | 43,17 ± 1,6 n=81 | |  |  |  |  |  |
| Figure | Labelling | Condition | Mean ± SEM | Source of Variation | F (DFn, DFd) | p value | Post-Hoc (Šídák) | p value | Test |
| Fig. S4A | VGLUT1+ | Singlets | 44,55 ± 4,57 n=6 | Interaction | F (2, 54) = 80,90 | ****p<0,0001 | Singlets - FASS |  | Two-way  MD ANOVA |
|  |  | FASS | 8,57 ± 1,53 n=14 | Condition | F (1, 54) = 0,0001234 | p=0,9912 | VGLUT1+ | ****p<0,0001 | |
|  |  |  |  | Staining | F (2, 54) = 293,7 | ****p<0,0001 | ++ | *p=0,0332 |  |
|  | ++ | Singlets | 1,79 ± 1,14 n=6 |  |  |  | EGFP+ | ****p<0,0001 | |
|  |  | FASS | 11,22 ± 1,38 n=14 | |  |  |  |  |  |
|  | EGFP+ | Singlets | 53,65 ± 4,29 n=6 | |  |  |  |  |  |
|  |  | FASS | 80,13 ± 2,27 n=14 | |  |  |  |  |  |
| Fig. S5E | Singlets | Singlets | 81,68 ± 0,8 n=10 | Interaction | F (2, 57) = 51,36 | ****p<0,0001 | Singlets - FASS |  | Two-way  MD ANOVA |
|  |  | FASS | 86,05 ± 1,47 n=11 | Condition | F (1, 57) = 1,450 | p=0,2335 | Singlets | p=0,6855 |  |
|  |  |  |  | Fluorescence | F (2, 57) = 156,0 | ****p<0,0001 | VENUS- | ****p<0,0001 | |
|  | VENUS- | Singlets | 52,71 ± 4,38 n=10 | |  |  | VENUS+ | ****p<0,0001 | |
|  |  | FASS | 23,85 ± 3,05 n=11 | |  |  |  |  |  |
|  | VENUS+ | Singlets | 18,48 ± 3,08 n=10 | |  |  |  |  |  |
|  |  | FASS | 52,06 ± 4,02 n=11 | |  |  |  |  |  |
| Fig. S6B | TH+ | Singlets | 36,51 ± 1,49 n=49 | Interaction | F (2, 255) = 251,5 | ****p<0,0001 | Singlets - FASS |  | Two-way  MD ANOVA |
|  |  | FASS | 66,65 ± 0,95 n=38 | Condition | F (1, 255) = 7,774e-006 | p=0,9978 | TH+ | ****p<0,0001 | |
|  |  |  |  | Staining | F (2, 255) = 297,4 | ****p<0,0001 | ++ | p=0,9576 |  |
|  | ++ | Singlets | 18,59 ± 1,51 n=49 | |  |  | Synaptopodin1+ | ****p<0,0001 | |
|  |  | FASS | 19,47 ± 0,84 n=38 | |  |  |  |  |  |
|  | Synaptopodin1+ | Singlets | 44,91 ± 1,72 n=49 | |  |  |  |  |  |
|  |  | FASS | 13,9 ± 0,58 n=38 | |  |  |  |  |  |
